# Aesthetic preference for art emerges from a weighted integration over hierarchically structured visual features in the brain

**DOI:** 10.1101/2020.02.09.940353

**Authors:** Kiyohito Iigaya, Sanghyun Yi, Iman A. Wahle, Koranis Tanwisuth, John P. O’Doherty

## Abstract

It is an open question whether preferences for visual art can be lawfully predicted from the basic constituent elements of a visual image. Moreover, little is known about how such preferences are actually constructed in the brain. Here we developed and tested a computational framework to gain an understanding of how the human brain constructs aesthetic value. We show that it is possible to explain human preferences for a piece of art based on an analysis of features present in the image. This was achieved by analyzing the visual properties of drawings and photographs by multiple means, ranging from image statistics extracted by computer vision tools, subjective human ratings about attributes, to a deep convolutional neural network. Crucially, it is possible to predict subjective value ratings not only within but also across individuals, speaking to the possibility that much of the variance in human visual preference is shared across individuals. Neuroimaging data revealed that preference computations occur in the brain by means of a graded hierarchical representation of lower and higher level features in the visual system. These features are in turn integrated to compute an overall subjective preference in the parietal and prefrontal cortex. Our findings suggest that rather than being idiosyncratic, human preferences for art can be explained at least in part as a product of a systematic neural integration over underlying visual features of an image. This work not only advances our understanding of the brain-wide computations underlying value construction but also brings new mechanistic insights to the study of visual aesthetics and art appreciation.

## Introduction

From ancient cave paintings to digital pictures posted on Instagram, the expression and appreciation of visual art can be found at the core of our human experience. As Kant famously pointed out, art is both subjective and universal.^1^ Each individual person may have his/her own taste, but a given piece of art can also appeal to a large number of people across cultures and history. This subjective universality raises a fundamental question: should artistic tastes be likened to the inscrutable, idiosyncratic, and irreducible, or is it possible to deduce lawful and generalizable principles by which humans form aesthetic opinions?

The nature of aesthetic judgment has long been subject to empirical investigation.^2–9^ Some studies have focused on the visual and psychological aspects of art might influence aesthetics (e.g., see^3, 5, 8, 10^), while other work has highlighted the brain regions whose activity level correlates with aesthetic values (e.g.,^11, 12^). However, attaining a mechanistic understanding of how the human brain computes aesthetic judgments in the first place from the raw visual input has thus far proved elusive.

A long-standing finding, which partly supports the idiosyncrasy of preference formation, is that prior experience with a specific stimulus can influence value judgment, such as the role of prior episodic memories involving the item, or prior associative history.^2–5, 13–15^ However, while the influence of past experience on current preference is undeniable, humans can express preferences for completely novel stimuli, suggesting that value judgments can be actively and dynamically computed.

Computationally, constructing a subjective value for a given piece of art is a non-trivial dimensionality reduction problem. The brain takes a massively high-dimensional input (e.g., a complex image presented to the retina) and, in the context of value judgments, eventually reduces this image to a one-dimensional scalar output (e.g., how much do I like this?). Performing dimensionality reduction at this scale is a fairly unconstrained problem; but nevertheless, the brain can generate a reliable preference rating for all kinds of visual inputs, raising the question of how exactly the brain accomplishes this computation.

In machine-learning, classification problems (e.g., dog vs. non-dog) are typically solved by projecting an input to a *feature space*.^16^ Each feature is a useful attribute that guides the classification of the input. Features can be engineered by taking easily observable characteristics of an object (e.g., its height and weight), or in other cases can be implicitly generated in a more abstracted and less easily interpretable manner (e.g., the activation patterns of hidden layers in deep artificial neural networks).

Previous studies support a feature-based framework as the base from which value judgments emerge in the brain. For example, one study^17^ implicated that subjective preference for food items can be expressed as a linear combination of various nutritive attributes, such as an item’s sugar and fat content. Another study suggested a similar coding of odor mixture components in food,^18^ where the values of individual order components, as well as the value of food as a whole, were tracked by orbitofrontal cortex (OFC). Further evidence supports attribute-based computations in prefrontal cortex (PFC), such as the distinction between whether a food is judged to be tasty or healthy,^19^ or even in judgments about the value of clothing,^20^ as well as the value of multi-attribute artificial stimuli (e.g., the movement and the color of dots) in humans and macaques.^21–23^

However, it is not clear if the same feature-based strategy used by the brain to derive value for items suited to a functional decomposition^24^ is also applied when performing the rather esoteric function of determining the aesthetic value of visual art. In previous work on feature integration, the features are natural and obvious properties of a stimulus such as a food’s fat content. However, in the case of visual imagery, the sheer complexity of the visual stimuli involved in one art piece, as well as the enormous variation between pieces, renders the task of identifying the relevant features that underpin this process exceedingly challenging. Even if relevant features are identified, it is unknown to what extent people may idiosyncratically select the features they use to shape their preferences and how they weigh those features to generate a value judgment.

Here, we aimed to establish a general mechanism that could underpin the construction of aesthetic preference. We first extracted features of an art image that have been theorized to play a role in aesthetic valuation.^8, 10, 25–27^ These features reflect subjective judgments about an image, and as such, we deemed them to be “high-level” features, as they required human judgment to determine their presence in an image. We augmented this with a bottom-up process that extracted visual features derived from each image’s statistics and visual properties, a feature set we labeled as “low-level”. We then used ratings from human participants’ across a large set of painting and photography images to ascertain the extent to which we could predict art preferences using our image feature set. Additionally, we applied a deep convolutional neural network (DCNN) to establish the degree to which features for computing visual preference might emerge spontaneously while processing visual images in an (approximately) brain-like architecture. Finally, we applied both linear and DCNN models to Functional magnetic resonance imaging (fMRI) data collected from human participants, which allowed us to identify the specific neural mechanisms underlying these feature representations during the evaluation of visual art, as well as to identify the mechanism by which such features are integrated to produce a value judgment.

## Results

### Linear feature summation (LFS) model predicts human valuation of visual art

Participants were asked to report how much they liked various pieces of art (images of paintings). The data were collected from both in-lab (N=7) and online participants using mechanical-turk (N=1359). On each trial, participants were presented with an image of a painting on a computer screen and asked to report how much they liked it on a scale of 1 (not at all) to 4 (very much) (Figure 1A). Each of the in-lab participants rated all of the paintings without repetition (1001 different paintings), while online participants rated approximately 60 stimuli, each drawn randomly from the image set. The stimulus set consisted of paintings from a broad range of art genres (Figure 1B), and each online participant saw images that were taken with equal proportions from different genres to avoid systematic biases related to style and time-period.

**Figure 1:**
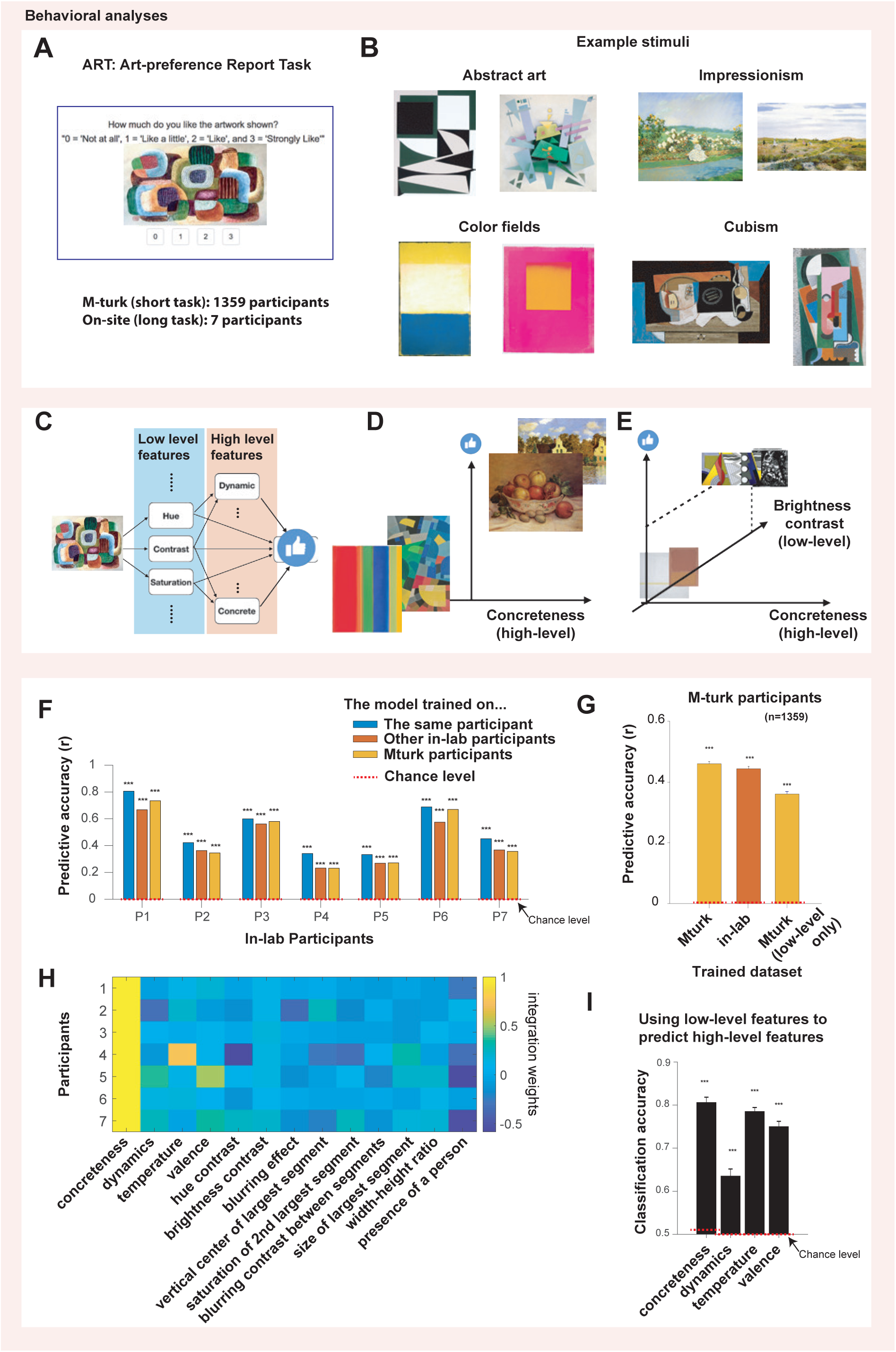
The linear feature summation (LFS) model was very successful in accounting for the subjective value of paintings. (**A**). The task (ART: art-liking rating task). Participants were asked to report how much they like a stimulus (a piece of artwork) shown on the screen using a four-point Likert rating ranging from 0 to 3. (**B**). Example stimuli. The images were taken from four categories from Wikiart.org.: Cubism, Impressionism, Abstract art and Color Fields, and supplemented with art stimuli previously used.^27^ Each m-turk participant performed approximately 60 trials, while in-lab participants performed 1001 trials (one trial per image). (**C**). Schematic of the LFS model. A visual stimulus (e.g., artwork) is decomposed into various low-level visual features (e.g., mean hue, mean contrast), as well as high-level features (e.g., concreteness, dynamics). We hypothesized that high-level features are constructed from low-level features, and that subjective value is constructed from a linear combination of all low and high-level features. (**D**). How features can help construct subjective value. In this example, preference was separated by the concreteness feature. (**E**). In this example, the value over the concreteness axis was the same for four images; but another feature, in this case, the brightness contrast, could separate preferences over art. (**F**). The LFS model with shared features captured in-lab participants’ art liking ratings. The predictive score, defined by the Pearson correlation coefficient between the model’s out-of-sample prediction and actual ratings, was significantly greater than chance for all subjects who performed the task in the lab. The model was trained on six participants and tested on the remaining participant (blue), trained and tested on the same participant (red), and trained on on-line participants and tested on in-lab participants (yellow). In-lab subjects performed a long task with 1001 trials. Statistical significance was tested against a null distribution of correlation scores constructed by the same analyses with permuted image labels. The chance level (the mean of the null distribution) is indicated by the dotted lines (at 0). The same set of features (shown in **H**) was used throughout the analysis. (**G**). Our model also successfully accounted for the on-line participants’ liking of the art stimuli. We trained the model on all-but-one participants and tested on the remaining participants (left). We also fit the model separately to in-lab participants and tested it independently on all on-line participants (middle). The model predicted liking ratings significantly in all cases, even when we used low-level attributes alone (right). Each on-line participant performed approximately 60 trials. The error bars show the mean and the SEM over participants. The chance level (the mean of the null distribution constructed in the same manner as F) is indicated by the dotted line. (**H**). Weights on shared features that were estimated for in-lab participants. We estimated weights by fitting individual participants separately. (**I**). The low-level features can predict the variance of high-level features. Classification accuracy (high or low values, split by medians) are shown. Note that though the prediction is highly significant, there is still a small amount of variance remaining that is unique to high-level features. The chance level (the mean of the null distribution) is indicated by the dotted line. The error bars indicate the standard errors over cross-validation partitions. In all panels, three stars indicate *p <* 0.001 against permutation tests.

Using this rating data, we tested our hypothesis that the subjective value of an individual painting can be constructed by integrating across features commonly shared across all paintings. For this, each image was decomposed into its fundamental visual and emotional features. These feature values are then integrated linearly, with each participant being assigned a unique set of features weights from which the model constructs a subjective preference (Figure 1C). This model embodies the notion that subjective values are computed in a feature space, whereby overall subjective value is computed as a weighted linear sum over feature content (Figure 1DE). We refer to this model as the Linear Feature Summation (LFS) model.

The LFS model extracts various low-level visual features from an input image using a combination of computer vision methods (e.g.,^25^). This approach computes numerical scores for different aspects of visual content in the image, such as the average hue and brightness of image segments, as well as the entirety of the image itself, as identified by machine learning techniques, e.g., Graph-Cuts^28^ (Details of this approach are described in the Methods section). Thus, we note that the LFS model performs a (weighted) linear sum over features, where features can be constructed non-linearly.

The LFS model also includes more abstract or “high-level” attributes that are likely to contribute to valuation. For this, we introduced three features based on previous studies^26, 27^ (the image is ‘abstract or concrete’, ‘dynamic or still’, ‘hot or cold’) as well as a fourth high-level feature concerning whether the image had a positive or negative emotional valence. Note that “valence” is not necessarily synonymous with valuation: if a piece of art denotes content with a negative emotional tone (e.g., Edvard Munch’s “The Scream”), it can still be judged to have a highly positive subjective value by the art appreciator. We hypothesized that these high-level features are constructed in downstream units using low-level features as input (Figure 1C). However, because we do not know the value of these high-level features a priori, following previous studies^26, 27^ we invited participants with familiarity and experience in art (n=13) to provide subjective judgments about the presence of each of these features in each of the images in our stimulus set (though we note a previous study found that artistic experience did not affect feature annotations^26^). We took the average score over these experts’ ratings as the input into the model representing the content of each high-level attribute feature for each image.

The final output of the model is a linear combination of low- and high-level features. We assumed that weights over the features are fixed for each individual, which is a necessary requirement to derive generalizable conclusions about the features used to generate valuation across images. As our high-level features were annotated by humans, we treat low-level and high-level features equally, in a non-hierarchical manner, in order to determine the overall predictive power of our LFS model.

We first determined a minimal set of features that can reliably capture rating scores across participants in order to gain insights into aesthetic preference universality. For this, we performed a group-level lasso regression on the data we collected in our in-depth in-lab study (n=7; each rated all 1001 images) using all of the low-level and high-level features that we constructed. By doing so, we removed from consideration those features that do not provide useful predictive information, ultimately selecting the features most uniquely predictive of subjective value, leaving 9 low-level and 4 high-level attribute features. The features include some low-level features computed from the entire images such as the ‘mean hue contrast’ and the ‘blurring effect’ as well as some low-level features computed using segmentation methods such as the ‘position and the size of the largest segment’, in addition to high-level features (please see the Methods for more details). Note that while the integration weights can be tuned for individual participant(s), the feature values for each image remain consistent for all participants.

We then asked how a linear regression model with these features can predict an individual’s liking for visual art. To our surprise, we found that we can predict subjective ratings in both a within-, and out-of-, participants manner; the model predicts subjective value not only when we trained the model’s weights on the same participant (using a cross-validated procedure) but also when we trained the weights on other in-lab participants, and even when we trained the weights in an entirely independent sample of online participants (Figure 1F). One potential concern may be that the model’s performance relies on the preferences for a particular art genre over the other (e.g., people may like impressionism over cubism); however, the same model trained on all images captures significant variations in preference *within* each art genre after taking out the effect of genre preferences (Figure S1), suggesting that the model captures variations in subjective preference both within and across genres.

Indeed, we found in a representation dissimilarity analysis over visual stimuli that low-level features seem to capture art genres, but high-level features go beyond the genres (Figure S2). We also found that we could reliably predict value ratings for online participants (Figure 1G), not only when training the model on the online participants’ data (using leave-one-out cross-validation) but also the model had been trained using in-lab participants’ data. We also tested the extent to which we can predict value from the low-level attributes alone. Removing the high-level features impaired predictive performance somewhat, but yielded highly significant prediction nonetheless (Figure 1G). These results suggest that a significant proportion of the variance in participants’ aesthetic ratings is universal, and can be generalized across people and art genres to a remarkable degree.

We then asked how the model’s integration weights for each participant can be varied across participants if we fit the model to each participant. Although we could predict each individual’s ratings by training the model on the ratings of others’, the degree to which each individual could be predicted from the pooled weights of other participants varied considerably. This suggests that while a common generic model of feature integration can predict individual liking ratings to a surprisingly high degree, there are also likely to be individual differences in how particular features are weighted, as we confirmed in the weights fit to each in-lab participant (Figure 1H). To identify potential clusters of individuals that might use features similarly, we turned to our much larger scale dataset of online participants. We fit our model’s weights to each online participant, and then fit a Gaussian mixture model to the estimated weights over participants. By comparing the Bayes Information Criteria score between models with a different number of Gaussians, we identified three clusters in the data (Figure S3). The majority of individuals (78%) in our online dataset were clustered according to the concreteness feature, apparently preferring images of scenery and impressionism. The remainder belong to one of two other groups: one (7%) with a strong preference for dynamic images (e.g., cubism), and the other (15%) had a large negative weight on concreteness and a positive weight on valence, exhibiting a preference for abstract art and color fields. We also noticed that the difference between clusters was not well described by art categories. As we noted, although we trained our model on the entire stimulus set, the model can still predict variation in preferences within each specific art genre (Figure S1).

The above results are based on a linear regression of low- and high-level features, but we also considered the likely possibility that high-level features are comprised of low-level features (illustrated in Figure 1C). To assess this, we probed the degree to which a linear combination of low-level features could predict the annotated ratings of high-level features. For this, we trained a linear support vector machine using all low-level features as input, and indeed we found that variance ascribed to high-level features could be predicted by low-level features (Figure 1I). This suggests that high-level features can be constructed using objective elements of the images, rather than subjective sensations, although the construction may well depend on additional nonlinear operations.

### The LFS model also predicts human valuation of photographs

One potential concern we had was that our ability to predict artwork rating scores using this linear model might be somehow idiosyncratic due to specific properties of the stimuli used in our stimulus-set. To address this, we investigated the extent to which our findings generalize to other kinds of visual images by using a new image database of 716 images;^29^ Figure 2A), this time involving photographs (as opposed to paintings) of various objects and scenes, including landscapes, animals, flowers, and pictures of food. We obtained ratings for these 716 images in a new m-Turk sample of 382 participants. Using the low-level attributes alone (these images were not annotated with high-level features), the linear integration model could reliably predict photograph ratings (Figure 2B). The model performed well when trained and tested on the photograph database, but to our surprise, the same model (as trained on photographs) could also predict the ratings for paintings that we collected in our first experiment, and vice versa (a model trained on the painting ratings could predict photograph ratings). Of note, accuracy was reduced if trained on paintings and tested on photographs (though still highly above chance), suggesting that the photographs enabled improved generalization (possibly because the set of photographs were more diverse). We stress that here, in all cases the model was trained and tested on completely separate sets of participants.

**Figure 2:**
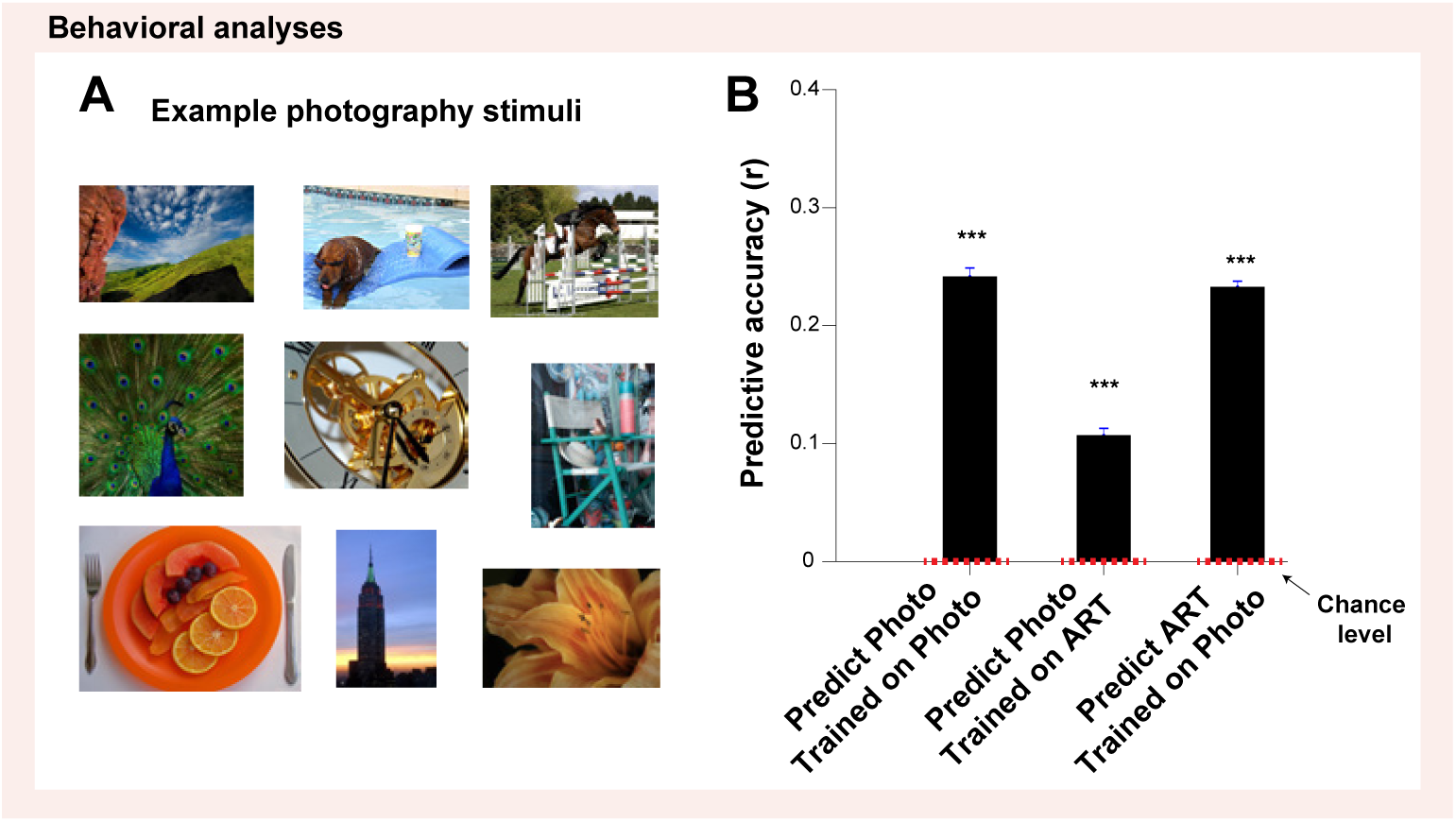
Our LFS model also predicts subjective liking ratings for various kinds of photographs. (**A**). Example stimuli from the photography dataset. We took a wide range of images from the online photography (AVA) dataset,^29^ and ran a further on-line experiment in new M-Turk participants (*n* = 382) to obtain value ratings for these images. (**B**). A linear model with low-level features alone captured liking ratings for photography. This model when trained on liking ratings for photography (from the current experiment) also captured liking ratings for paintings (from the previous experiment described in Figure 1), and the model trained on liking ratings for paintings could also liking ratings for photography. We note that in all cases the model was trained and tested on completely separate sets of participants. The significance was tested against the null distribution constructed from the analysis with permuted image labels. The error bars indicate the mean and the SEM.

### A deep convolutional neural network (DCNN) model predicts human liking ratings for visual art

We now have shown that our LFS model can capture subjective preference for visual art; however, as we engineered the model’s features using a mixture of prior literature and bottom-up machine learning tools, we do not know if this strategy has any biological import. To investigate this further, we next asked whether the LFS model’s feature representations can spontaneously emerge in a neural network that is trained to predict subjective preference in the absence of feature engineering. For this, we utilized a standard deep convolutional neural network (DCNN; VGG 16^30^), that had been pre-trained for object recognition with ImageNet.^31^ We used this network with fixed pre-trained weights in convolutional layers, but trained the weights for the last three fully-connected layers on averaged liking ratings. Mirroring the results of our LFS model, we found that our DCNN model can predict human participants’ liking ratings across all participants (Figure 3A). This shows that it is indeed possible to predict preferences for visual art using a deep-learning approach without explicitly selecting stimulus features. In a supplementary analysis, we also opened the convolutional layers to training, but saw no improvement.

**Figure 3:**
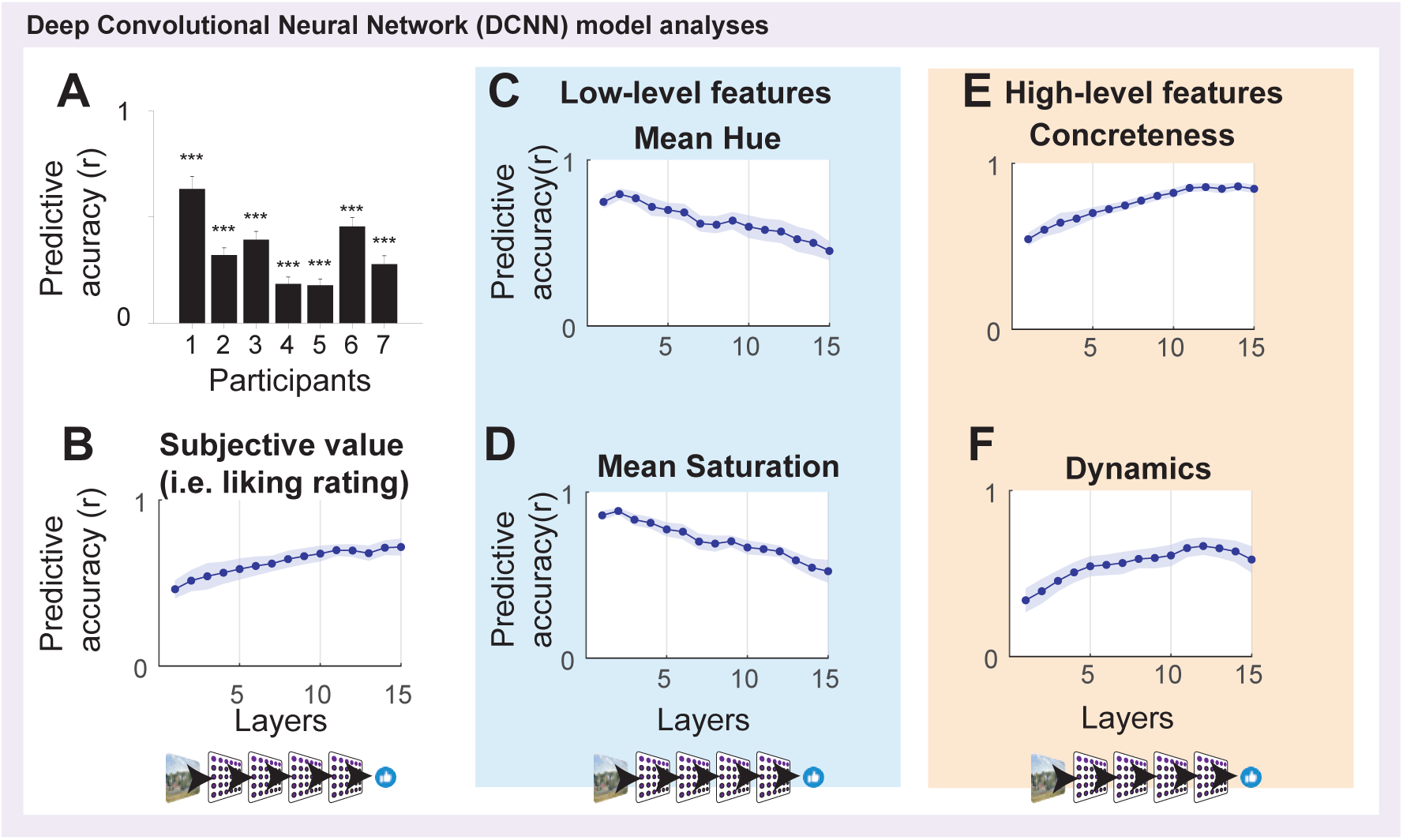
A deep convolutional neural network (DCNN) can predict subjective values (i.e., liking ratings) of art stimuli, and the features that we introduced to our LFS model spontaneously emerge in the hidden layers of the network. We utilized a standard convolutional neural network (VGG 16^30^) that came pre-trained on object recognition with ImageNet,^31^ consisting of 13 convolutional and three fully connected layers. We trained the last three fully connected layers of our network on average art liking scores without explicitly teaching the network about the LFS model’s features. (**A**).The neural network could successfully predict human participants’ liking ratings significantly greater than chance across all participants. The significance (*p <* 0.001, indicated by three stars) was tested by a permutation test. (**B**). We found that we can decode average liking ratings using activation patterns in each of the hidden layers. The predictive accuracy was defined by the Pearson correlation between (out-of- sample) model’s predictions and the data. For this, we used a (ridge) linear regression to predict liking ratings from each hidden layer. We first reduced the dimensions of each layer with a PCA, taking top PCs that capture 80% of the variance in each layer. The accuracy gradually increases over layers despite the fact that most layers (layers 1-13) were not trained on liking ratings but on ImageNet classifications alone. (**C,D**). When performing the same analysis with the LFS model’s features, we found some low-level visual features with significantly decreasing predictive accuracy over hidden layers (e.g., the mean hue and the mean saturation). We also found that a few computationally demanding low-level features showed the opposite trend (see the main text). (**E,F**). We found some high-level visual features show significantly increasing predictive accuracy over hidden layers (e.g., concreteness and dynamics). We also found that temperature, which we introduced as a putative high-level feature, actually shows the opposite trend, likely because it is a color-based feature that can be straightforwardly computed from pixel data.

### The LFS model’s features emerge spontaneously in the DCNN model’s hidden layers

We turn now to ask whether the features used in our LFS model are spontaneously encoded in the neural network. Mirroring our illustration of the LFS model in Figure 1C, we hypothe-sized that low-level visual features would be represented in early layers of the DCNN, while more abstract high-level attributes would be represented in later layers of the network. For our investigation, we performed decoding analyses to predict high- and low-level feature values using the activation patterns in each hidden layer. We first reduced the dimensions of each layer using principal component analysis (PCA), and using the top principal components (PCs) that capture 80% of the variance of each layer, we trained a regression model for a given variable that we aimed to predict (e.g., ratings or a feature) using the PCs of each layer.

We first tested to see if we can predict subjective liking ratings using the hidden layer activation patterns. We were able to decode subjective ratings across all layers, but noted that decodability gradually increased for layers deeper in the network (Figure 3B). This came as a surprise since all but the last three layers (layers 1 to 13, out of 16) were pre-trained not on the rating scales being decoded, but on image classifications alone using ImageNet, hinting at a tight relationship between value coding and image recognition.

We then tested to see how the hidden layers related to the LFS model’s features. This analysis showed that hidden layers could predict all 23 features included in the LFS model. Consistent with our hypothesis, six (of the 19) putative low-level features tested were represented more robustly in early layers, as shown by a significantly negative decoding slope across layers (Figure 3CD). We also found for four (out of 19) low-level features had a decoding accuracy that increased as a function of the depth of the layer, suggesting those low-level features, in fact, may be better identified as high-level features. However, we note that the overall predictive accuracy of these features was low compared to those showing negative slopes. These positive slope features include: “the presence of a person”, “the mass center for the largest segment”, “mass variance of the largest segment”, and “entropy in the 2nd largest segment”, all of which require relatively complex computations (e.g., segmentation and the identification of the location of the segments) compared to the ones showing negative slopes (e.g., average saturation). We note that this result is consistent with a previous electrophysiological and computational modeling study in macaques,^32^ which reported that the position of an object on the screen is more robustly represented in higher visual areas and deeper layers, as position identifications of a segment and an object likely involve similar computations. These object-related features were also referred to as ‘low-level’ features,^32^ in line with our original reference. The other 9 low-level features tested did not show either a strong positive or negative slope.

Similarly, for the putative high-level features, we found that two (of 4) features were more robustly represented in later layers (Figure 3EF). However, “temperature”, which as labelled as a high-level feature, showed a significant negative slope. Given that this feature was based on color palettes in the image (i.e., whether the color palette is hot or cold), this feature’s variance may already be well captured by low-level image statistics. The fourth putative high-level feature, valence, did not show either an increasing or decreasing trend in decoding across layers. Thus, the DCNN allows a more principled means to identify low and high-level features, enabling us to label 6 features as low-level (based on greater representation of those features in earlier layers of the network), and 6 as high-level features, indicated by representations that are present to a greater extent in later layers of the network. These LFS model-based analyses on DCNN sheds light onto what are often-considered-to-be “black box” computations in deep artificial neural networks, and may provide an empirical definition of computational complexity in feature extraction of visual as well as other sensory inputs.

Taken together, our DCNN analyses suggest that our conceptualized LFS model (Figure 1C) is, in fact, a natural consequence of training the neural network on object recognition and predicting subjective aesthetic value, without requiring any explicit feature engineering.

### The subjective value of art is represented in the medial prefrontal cortex (mPFC)

We have shown that a simple linear feature summation model (LFS) can predict the subjective valuation of visual art and that those features are naturally encoded in a deep convolutional neural network that itself can successfully predict art valuation. A remaining question is whether and how the actual human brain performs such a feature-based computation when making subjective value judgments about art. Therefore, we next ran an fMRI study in which we aimed to link our two computational models (the LFS model and the DCNN model) to neural data as a test of how feature-based value construction may be realized in the brain. Rather than scan a large number of participants for a short period of time to obtain group averaged results, here, we engaged in deep fMRI scanning of a smaller group of individuals (n=6), who each completed 1000 trials of our art rating task each over four days of scanning (Figure 4A). This allowed us to test for the representation of the features in each individual participant with sufficient fidelity to perform reliable single subject inference, an approach typically used in the visual neuroscience literature (e.g.,^33, 34^).

**Figure 4:**
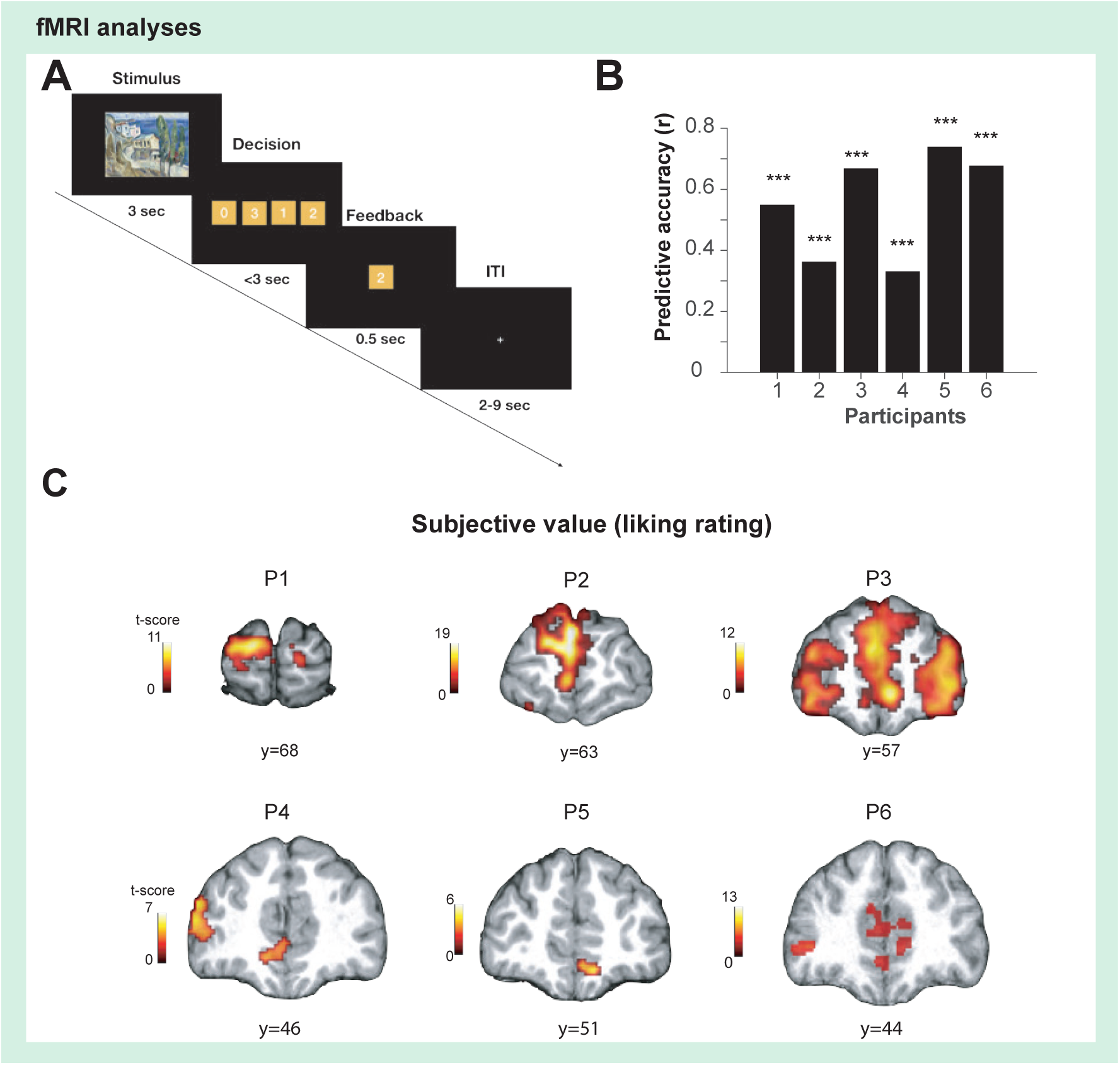
Neuroimaging experiments. (**A**). We administered our task (ART) to human participants in an fMRI experiment. Each participant completed 20 scan sessions spread over four separate days (1,000 trials in total with no repetition of the same stimuli). On each trial, a participant was presented with a visual art stimulus (paintings) for 3 sec. The art stimuli were the same as what we used in our behavioral study. After the stimulus presentation, a participant was presented with a set of possible ratings (0,1,2,3), where they had to choose one option within 3 seconds, followed by brief feedback with their selected rating (0.5 sec). The positions of the numbers were randomized across trials, and the order of presented stimuli was randomized across participants. (**B**). The LFS model successfully predicts participants’ liking ratings for the art stimuli. The model was fit to each participant (cross-validated). The significance was tested by the permutation test. Three stars indicate *p <* 0.001. (**C**). Subjective value (i.e., liking rating). Subjective value for art stimuli at the time of stimulus onset was found in the medial prefrontal cortex in all six fMRI participants (whole-brain cFWE *p <* 0.05 with height threshold at *p <* 0.001).

We first identified a shared feature set for our fMRI analyses using behavioral data pooled across all six fMRI and seven behavior-only in-lab participants. Since our interests hinge on the representational relationship between low- and high-level features in the brain, we first identified a set of low-level features that predict behavioral liking ratings with the same procedure as before, and further augmented this to a full feature set by including the four human-annotated features. By doing so, we aimed to identify signals that are uniquely correlated with low-level and high-level features (i.e., partial correlations between features and fMRI signals). In the following, we present the results with our original reference to features (four human-annotated features as high-level features and the rest as low-level features), but we also confirmed in our supplementary analyses that referring to features according to our DCNN analysis, i.e., by using the slopes of our decoding results, did not change the results of our fMRI analyses qualitatively and does not affect our conclusions (see Figure S7).

Using the shared feature set, we fit the LFS model to each fMRI participant, confirming that the model could predict subjective art ratings across participants, once again replicating our previous behavioral findings (Figure 4B; see Figures S4 and S5 for the estimated weights for each participants and their correlations).

We then examined for brain regions whose activation patterns correlated with the subjective liking ratings of each individual stimulus at the time of stimulus onset. Confirming an extensive prior literature (e.g.,^11, 12, 35–39)^, we found that voxels in the medial prefrontal cortex (mPFC) were positively correlated with subjective value across participants (Figure 4C;see also Figure S6 for other regions).

### Visual stream shows input (visual art) transformation of low- to high-level features

As illustrated in Figure 1C, and reflecting our hypothesis regarding the encoding of low vs. high-level features across layers of the DCNN, we hypothesized that the brain would decompose visual input similarly, with early visual regions first tracking low-level features, and with downstream regions constructing high-level features. Specifically, we analyzed visual cortical regions in the ventral and dorsal visual stream^41^ to test the degree to which low-level and high-level features are encoded in a graded, hierarchical manner. In pursuit of this, we constructed a GLM that included the shared feature time-locked to stimulus onset. We identified voxels that are significantly modulated by at least one low-level feature by performing an F-test over the low-level feature beta estimates, repeating the same analysis with high-level features. We then compared the proportion of voxels that were significantly correlated with low-level features vs. high-level features in each region of interest in both the ventral and dorsal visual streams. Regions of interest were independently identified by means of a detailed probabilistic visual topographical map.^41^ Consistent with our hypothesis, our findings suggest that the low- and high-level features are indeed represented in the visual stream in a graded hierarchical manner. Namely, the relative encoding of high-level features with respect to low-level features dramatically increases across the visual ventral stream (Figure 5A). We found a similar, hierarchical organization in the dorsolateral visual stream (Figure 5B), albeit less clearly demarcated than in the ventral case (a similar pattern of results was obtained if we used the feature labels identified from the DCNN decoding analysis (Figure S7)).

**Figure 5:**
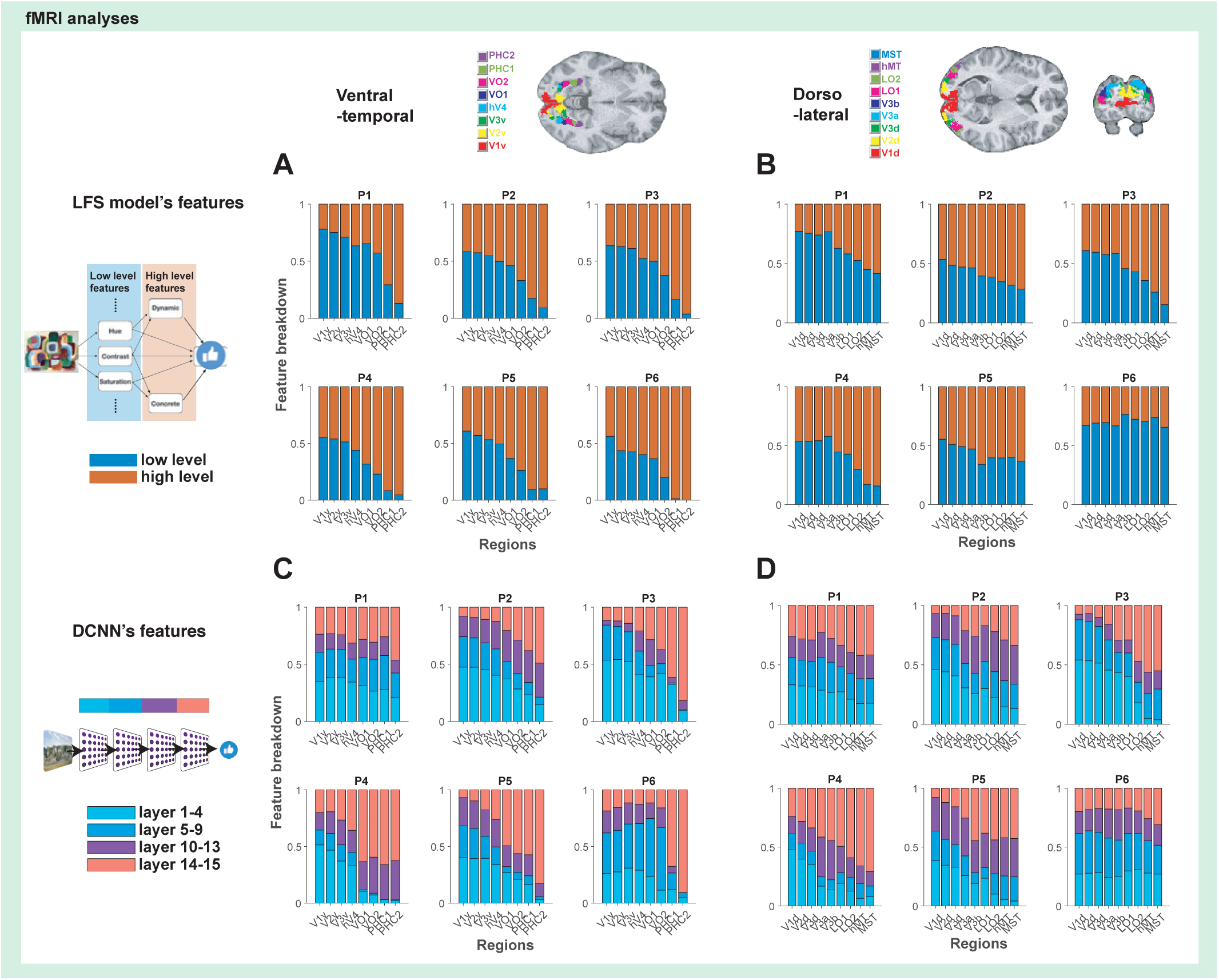
fMRI signals in visual cortical regions show similarity to our LFS model and DCNN model. (**A**). Encoding of low and high-level features in the visual ventral-temporal stream in a graded hierarchical manner. In general, the relative encoding of high-level features with respect to low-level features increases dramatically across the ventral-temporal stream. To illustrate the anatomical location of each ROI, the maximum probabilistic map^40^ is shown color-coded on the structural MR image at the top. The proportion of voxels that significantly correlated with low-level features (blue; F-test *p <* 0.001) against high-level features (red; F-test *p <* 0.001) are shown for each ROI. Please see the Methods section for detail. (**B**). Encoding low and high-level features in the dorsolateral visual stream. The anatomical location of each ROI^40^ is color-coded on the structural MR image. (**C**). Encoding of DCNN features (hidden layers’ activation patterns) in the ventral-temporal stream. The top three principal components from each layer of the DCNN were used as features in this analysis. In general, early regions more heavily encode representations found in early layers of the DCNN while higher-order regions encode representations found in deeper CNN layers. The proportion of voxels that significantly correlated with PCs of convolutional layers 1 to 4 (light blue), convolutional layers 5 to 9 (blue), convolutional layers 10 to 13 (purple), fully connected layers 14-15 (pink) are shown for each ROI. The significance was set at *p <* 0.001 by F-test. (**D**). Encoding of DCNN features in the dorsolateral visual stream.

**Figure 6:**
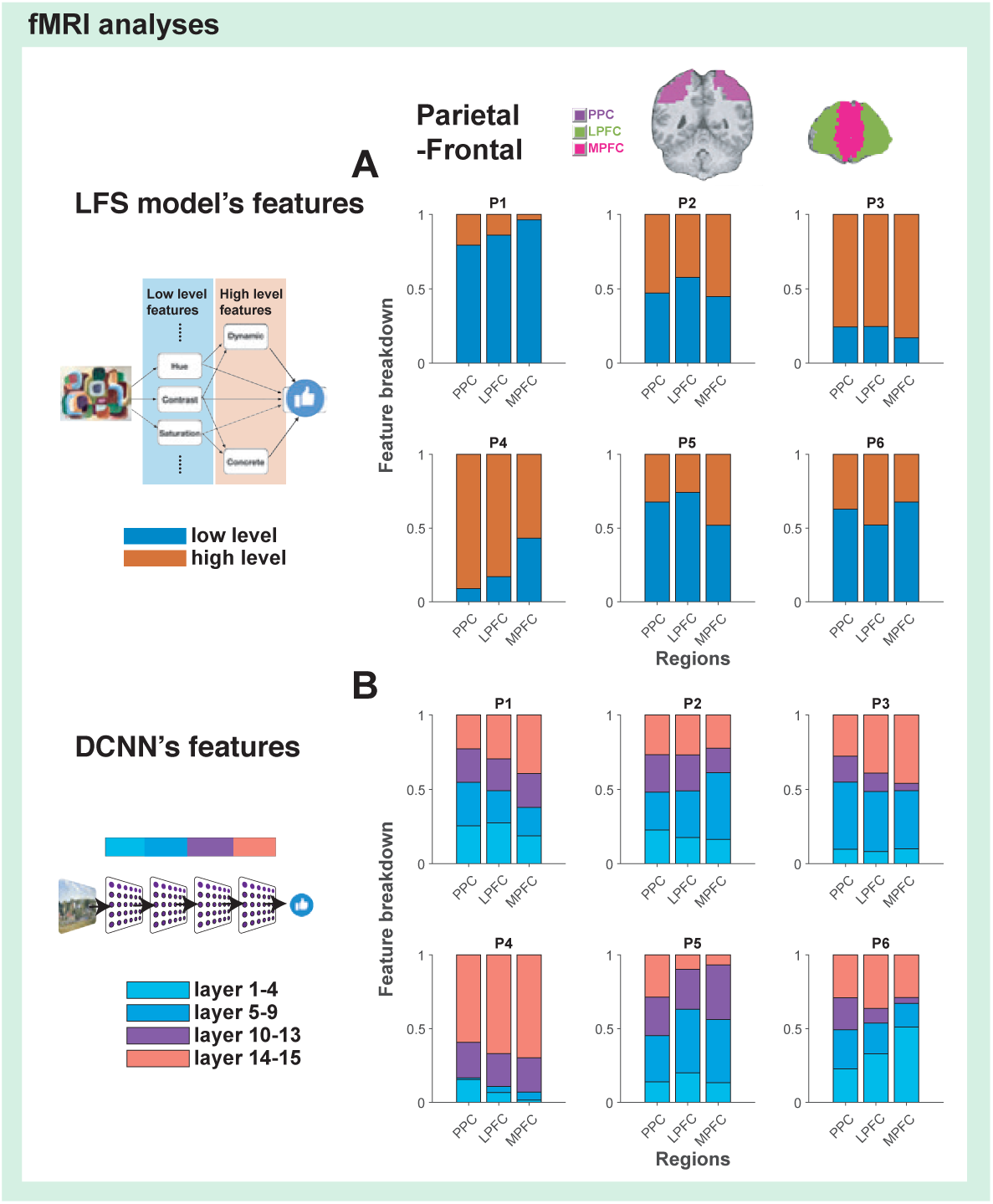
Parietal and prefrontal cortex encode features in a mixed manner. (**A**). Encoding of low- and high-level features from the LFS model in posterior parietal cortex (PPC), lateral prefrontal cortex (lPFC) and medial prefrontal cortex (mPFC). The ROIs used in this analysis are indicated by colors shown in a structural MR image at the top. (**B**). Encoding of the DCNN features (activation patterns in the hidden layers) in PPC and PFC. The same analysis method as Figure 5 was used.

We then tested whether activity patterns in these regions resembles the computations performed by the hidden layers of the DCNN model. We extracted the first three principal components from each layer of the DCNN, and included each as regressors in a GLM. Indeed, we found evidence that both the ventral and dorsal visual stream exhibits a similar hierarchical organization to that of the DCNN, such that lower visual areas correlated better with activity in the lower levels of the DCNN, while higher-order visual areas (in both visual streams) tend to correlate better with activity in deeper layers of the DCNN (Figure 5C, D).

### PPC and PFC show mixed coding of low- and high-level features

Building on this identification of the features’ representational structure, we asked how these representations are projected to downstream regions of association cortex^42, 43^ to play a role in stimulus valuation. We performed the same analysis with the same GLM as before in regions of interest that included the posterior parietal cortex (PPC), lateral prefrontal cortex (lPFC) and medial prefrontal cortex (mPFC). We found that both the LFS model features and the DCNN’s layers were represented in these regions in a mixed manner.^44, 45^ However, probing these regions showed no clear evidence for a progression of the hierarchical representations that we had observed in visual cortex; instead, each of these regions appeared to represent both low and high-level features to a similar degree. This suggests that, as we will see, these regions appear to play a primary role in feature integration as required for subjective value computations.

### Features encoded in PPC and lPFC are strongly coupled to the subjective value of visual art in mPFC

Having established that both the engineered LFS model and the emergent DCNN’s model features are hierarchically represented in the brain, we asked if and how these features are ultimately integrated to compute the subjective value of visual art. First, we analyzed how aesthetic value is represented across cortical regions alongside the model’s features by adding the participant’s subjective ratings to the GLM. We found that subjective values are, in general, more strongly represented in the PPC as well as in the lateral and medial PFC than in early and late visual areas (Figures 7A and S8). Furthermore, value signals appeared to become more prominent in medial prefrontal cortex compared to the lateral parietal and prefrontal regions (consistent with a large prior literature, e.g.,^11, 35–39, 46^). As further validation of our earlier feature encoding analyses, we found that the pattern of a hierarchical representation of features found in visual regions was unaltered by the inclusion of ratings in the GLM (Figure S9).

**Figure 7:**
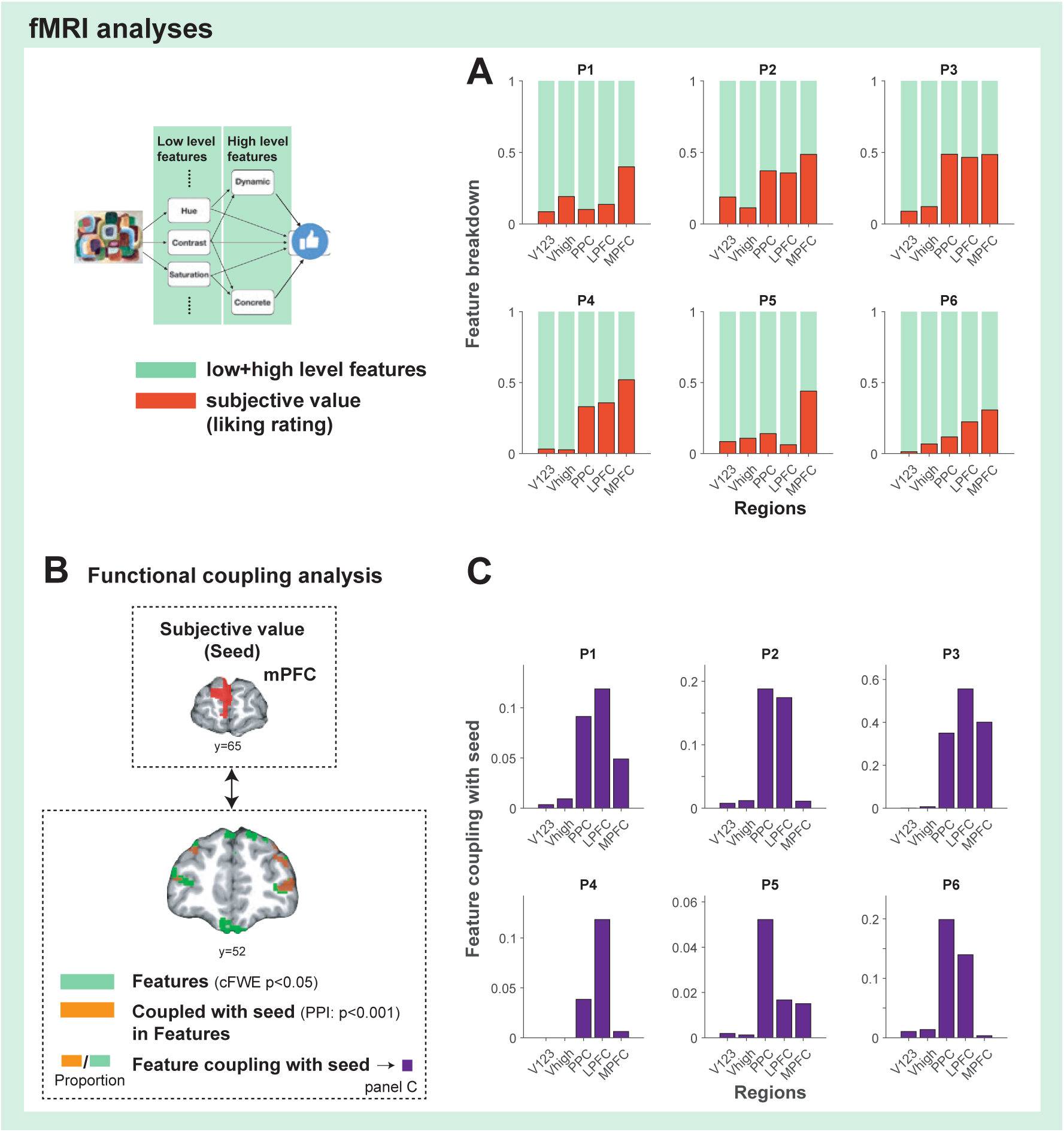
Features represented in PPC and lateral PFC are integrated to medial PFC when constructing the subjective value of visual art. (**A**). Encoding of low and high level features (green) and liking ratings (red) across brain regions. Note that the ROIs for the visual areas are now grouped as V1-2-3 (V1, V2 and V3) and V-high (Visual areas higher than V3). Please see the Methods section for detail. (**B**). Functional coupling analysis to test how feature representations are coupled with subjective value. We identified regions that encode features (green), by performing an F-test (*p <* 0.05 whole-brain cFWE with the height threshold *p <* 0.001). We also performed a psychophysiological interaction (PPI) analysis (orange: *p <* 0.001 uncorrected) to determine the regions that are coupled to the seed regions in mPFC that encode subjective value (i.e., liking rating) while stimulus presentation (red: seed, please see Figure S10). We then asked what is the proportion that these two overlap in feature encoding voxels in a given ROI. (**D**). The results of the functional coupling analysis show that features that are represented in the PPC and lPFC are coupled with the region in mPFC encoding subjective value. This result dramatically contrasts with our control analysis focusing on ITI instead of stimulus presentations (Figure S12).

These results suggest that the representations present in the PPC and lateral PFC are leveraged to construct subjective values in mPFC. To test this, we performed a psychological-physiological interaction (PPI) analysis, examining which of the voxels that represent the LFS model’s features are coupled with regions that represent subjective value (Figure 7B **and** S10). We then tested for the fraction of feature-encoding voxels that are also correlated with the PPI regressor across each ROI. This analysis revealed that the overlap was most prominent in the PPC and lPFC, while there was virtually no overlap in the visual areas at all (Figure 7C). More detailed decomposition of the PFC ROI from the same analysis shows the contribution of individual sub-regions of lateral and medial PFC (Figure S11). We also performed a control PPI analysis to test the specificity of the coupling to an experimental epoch by constructing a similar PPI regressor locked to the epoch of inter-trial-intervals (ITIs). This analysis showed a dramatically altered coupling that did not involve the same PPC and PFC regions (Figure S12). These findings suggest that both PPC and LPFC are specifically involved in the integration of feature representations in order to compute the subjective value, and that this integration depends on connections between feature representations in PPC/lFPC and subjective value representations in mPFC.

## Discussion

Whether we can lawfully account for personal preferences in the aesthetic appreciation of art has long been an open question in the arts and sciences.^1, 2, 4, 6^ Here, we addressed this question by engineering a hierarchical linear feature summation (LFS) model that generates subjective preference according to a weighted mixture of explicitly designed stimulus features. This model was verified with large-scale behavioral experiments, and contrasted to a deep convolutional neural network (DCNN) model as well as in in-depth focused, within-subject, neuroimaging experiments. We found that it is indeed possible to predict subjective valuations of both paintings and photography, and we demonstrate how the brain transforms visual stimuli into subjective value.

Our results indicate that linearly integrating a small set of visual features can explain human preferences for paintings and photography. Not only is it possible to predict an individual’s ratings based on that particular individual’s prior ratings for other images, but we also found that this strategy allowed us to predict one individual’s preferences from the preferences of others, even for novel stimuli. This is achievable likely because the majority of participants have similar preferences, and the model efficiently extracted the tastes, as shown by our clustering analysis whereby one dominant cluster was found to account for the majority of participants’ liking ratings (Figure S3).

Here, utilizing a set of interpretable visual and emotional features, we showed that these features are employed by the human brain to make value judgments for art. We note that this is by no means a complete enumeration of the features used by the brain. For instance, the semantic meaning of a painting, its historical importance, as well as memories of past experiences elicited by the painting, are also likely to play important roles (see, e.g.,^5, 8, 47^). Thus, rather than offering a feature catalogue, our findings shed light on the general principles by which feature integration yields to aesthetic valuation. However, the features that we identified are likely to be important, particularly as we utilized a reasonably large set of potential features in our initial feature set which was subsequently narrowed down to a set of only the most relevant features.

We found evidence using a DCNN model that the features engineered for use in the LFS model spontaneously emerge in a neurally plausible manner. Our deep network model was not explicitly trained on any of the LFS model’s features, nor were the convolutional hidden layers of the network trained on liking ratings (only the later fully connected layers were trained using rating scores). Nevertheless, we were able to identify LFS features from the hidden layers of the network, which suggests that the features used for aesthetic valuation likely emerge spontaneously through more basic and generalizable aspects of visual processing. Further, those features may well be utilized for a wide range of visual tasks, including object classification, prediction, and identification. Thus, we speculate that these findings suggesting a common feature space shared across different tasks may provide insights into transfer learning^48^ in machine learning.

One important consideration is whether linear feature operations are sufficient to describe the computations underlying aesthetic valuation. Notably, the highly non-linear deep network did not substantively outperform the simple linear model. However, in the LFS model, the feature extraction process itself is not necessarily linear (e.g., segmentation). As such, our results do not rule out the possibility of nonlinearity in feature extraction processes in the brain, but they do suggest that the final feature value integration for computing subjective art valuation can be approximated by a linear operation. This computational scheme resonates with a widely-used machine-learning technique referred to as the kernel method, whereby inputs are transformed into a high-dimensional feature space in which categories are linearly separable,^49, 50^ as well as with high-dimensional task-related variables represented in PFC.^44^

We obtained in-depth fMRI data from participants who underwent the art rating task as a means of investigating how aesthetic value computation is actually implemented in the brain. Focusing first on the visual system, we found that low-level features were represented more robustly in early visual cortical areas, while high-level features were increasingly represented in higher-order visual areas. These results support a hierarchical representation of the features required for valuation of visual imagery, and further support a model whereby lower-level features extracted by early visual regions are integrated to produce higher-level features in the higher visual system.^10^ We note that the notion of hierarchical representations in the visual system is well established in the domain of object recognition,^51–54^ where similarities between the computations in different layers of feed-forward deep neural networks trained on visual object recognition and neural activity in primate and human visual cortex has been extensively documented (e.g.,^32, 55–57)^. Our results substantially extend these findings by showing that features relevant to a very different behavioral task – forming value judgments, are also represented robustly in a similar hierarchical fashion. Crucially, the flexibility afforded by a feature-based mechanism of value construction ensures that value judgments can be formed even for stimuli that have never before been seen, or in circumstances where the goal of valuation varies (e.g., selecting a piece of art as a gift).

Our findings suggest that the processes through which feature representations are transformed and mapped into a singular subjective value dimension occur in a network of brain regions, including the posterior parietal cortex (PPC), lateral and medial frontal cortices (lPFC ad mPFC). We show that features relevant for computing subjective value are widely represented in lPFC and PPC, whereas subjective value signals are more robustly represented in parietal and frontal regions, with the strongest representation in mPFC. Further, lPFC and PPC regions encoding low- and high-level features enhanced their coupling with mPFC encoding subjective value at the time of image presentation. Although we cannot make direct inferences about the directionality of the connectivity effects, these findings are compatible with a framework in which low and high-level feature representations in lPFC and PPC are utilized to construct value representations in mPFC. This possibility is further supported by our analysis showing a similarity between representations in the hidden layers of the DCNN and fMRI responses throughout visual as well as association cortices. Similar to the linear feature based analysis, we found evidence to support a significant correlation between the DCNN’s representations of early layers and fMRI responses in the early visual system, while later layers of the DCNN were increasingly correlated with activity in higher-order visual areas. Taken together, these findings are consistent with a large-scale processing hierarchy in the brain that extends from early visual cortex to medial prefrontal cortex, whereby visual inputs are transformed into various features through the visual stream. These features are then projected to PPC and lPFC, and subsequently integrated into subjective value judgment in mPFC.

The present findings offer a mechanism through which artistic preferences can be predicted. It is of course important to note that aesthetic experience more broadly defined goes beyond the simple one-dimensional liking rating (a proxy of valuation) that we study here (e.g.,^4, 6, 8, 47^), and that judgments can be context-dependent.^58^ Art is likely to be perceived along many dimensions, of which valuation is but one, with some dimensions relying more on idiosyncratic experience than others. Nevertheless, we speculate that just as it is possible to explain aesthetic valuation in terms of underlying features, many other aspects of the experience of art can also likely be decomposed into more atomic feature-based computations, with different dimensions employing different weights over those features. Indeed, subjective value can itself be considered to be a “feature” in a feature space, albeit a high-level one, alongside other judgments that might be made about a piece of art. However, given its centrality in shaping and motivating behavior in humans and other animals, we would argue that subjective value is nevertheless a very fundamental aspect of the evaluation of any stimulus, whether it be a piece of art or otherwise.

To conclude, our findings demonstrate that once relevant features are extracted from a visual image, it is possible to dynamically construct values that can account for actual art preferences. In the brain, the active construction of value appears to involve neural circuits associated with feature representation in the parietal and prefrontal cortex. Thus, far from being inscrutable and idiosyncratic, with modern computational and machine-learning tools, aesthetic valuation can be lawfully described and its mechanisms illuminated at both computational and neural levels.

## Methods

### Participants

#### Online study

A total of 1936 volunteers (female: 883 (45.6%). age 18-24 yr: 285 (14.8%); 25-34 yr: 823 (42.8%); 35-44 yr: 435 (22.6%); 45 yr and above: 382 (19.8%)) participated in our on-line studies in the Amazon Mechanical Turk (M-turk). 1545 of them participated in the ART task, and 391 of them participated in the AVA photo task. Among these, participants who missed trials and failed to complete 50 trials were excluded from our analyses, leaving us with online 1359 participants in the ART task data and 382 participants in the AVA photo task data.

#### In-lab study

Thirteen volunteers (female: 9. age 18-24 yr: 5; 25-34 yr: 5; 35-44 yr: 3) were recruited to our in-lab studies, where seven of them participated in our behavioral task, and six of them participated in our fMRI task.

Additionally, thirteen art-experienced participants (female: 6. age 18-24 yr: 3; 25-34 yr: 9; 35-44 yr: 1) were invited to evaluate the high-level feature values. These participants for annotation were primarily recruited from the ArtCenter College of Design community. Participants provided informed consent for their participation in the study, which was approved by the Caltech IRB.

### Stimuli

Painting stimuli were taken from the visual art encyclopedia www.wikiart.org. Using a script that randomly selects images in a given category of art, we downloaded 206 or 207 images from four categories of art (825 in total). The categories were ‘Abstract Art’, ‘Impressionism’, ‘Color Fields’, and ‘Cubism’. We randomly downloaded images with each tag using our custom code in order to avoid subjective bias. For the in-lab and fMRI (but not on-line) studies, we supplemented this database with an additional 176 paintings that were used in a previous study.^27^ For the fMRI study, one image was excluded from the full set of 1001 images to have an equal number of trials per run (50 images*/*run 20 × runs = 1000 images).

Picture images were taken from the Aesthetic Visual Analysis (AVA) dataset. This dataset consists of images from multiple online photo contests. We took images from the following categories (about 90 images from each): ‘Animals’, ‘Floral’, ‘Nature’, ‘Sky’, ‘Still Life’,‘Advertisement’, ‘Sky’, and ‘Abstract Pictures’. In a total of 716 images were used.

## Tasks

### Behavioural task

#### Liking rating task

On each trial, participants were presented with an image of the artwork (in the Art-liking Rating Task: ART) or a picture image (in the AVA photo task) on the computer screen. Participants reported within 6 seconds how much they like the artwork (or the picture image), by pressing buttons corresponding to a scale that ranged from 0, 1, 2, 3, where 0 = not like at all, 1= like a little, 2 = like, and 3 = strongly like, presented at the mottom of the image. Each of the on-line ART participants performed on average 57 trials of the rating task, followed by a familiarity task in which they reported if they could recognize the name of the artist who painted the artwork for the same images that they reported their liking ratings. The images for each online participant were drawn to balance different art genres. On-site ART participants performed 1001 trials of rating tasks. On-site participants had a chance to take a short break approximately every 100 trials. Each of the on-line AVA photo task participants performed on average 115 trials of the rating task.

#### Feature annotation

The four high-level features were annotated in a manner following.^26, 27^ On each trial, participants were asked about the feature value of a given stimulus, ranged from −2, −1, 0, 1, 2. Following,^26^ example figures showing extreme feature values are always shown on the screen as a reference (please see Figure S13). Each participants completed four separate tasks (for four features) in a random order, where each task consists of 1001 trials (with 1001 images).

### fMRI task

On each trial, participants were presented with an image of the artwork on the computer screen for three seconds. Participants were then presented with a scale from 0, 1, 2, 3 in which they had to indicate how much they liked the artwork. The location of each numerical score was randomized across trials. Participants had to press a button of a button box that they hold with both hands to indicate their rating within three seconds, where each of four buttons corresponded to a particular location on the screen from left to right. The left (right) two buttons were instructed to be pressed by their left (right) thumb. After a brief feedback period showing their chosen rating (0.5 sec), a center cross was shown for inter-trial intervals (jittered between 2 to 9 seconds). Each run consists of 50 trials. Participants were invited to the study over four days to complete twenty runs, where participants completed on average five runs on each day.

### fMRI data acquisition

fMRI data were acquired on a Siemens Prisma 3T scanner at the Caltech Brain Imaging Center (Pasadena, CA). With a 32-channel radiofrequency coil, a multi-band echo-planar imaging (EPI) sequence was employed with the following parameters: 72 axial slices (whole-brain), A-P phase encoding, −30 degrees slice orientation from AC-PC line, echo time (TE) of 30ms, multi-band acceleration of 4, repetition time (TR) of 1.12s, 54-degree flip angle, 2mm isotropic resolution, echo spacing of 0.56ms. 192mm x 192mm field of view.

Positive and negative polarity EPI-based fieldmaps were collected before each run with very similar factors as the functional sequence described above (same acquisition box, number of slices, resolution, echo spacing, bandwidth and EPI factor), single band, TE of 50ms, TR of 5.13s, 90-degree flip angle.

T1-weighted and T2-weighted structural images were also acquired once for each participant with 0.9mm isotropic resolution.

### fMRI data processing

Results included in this manuscript come from preprocessing performed using *fMRIPrep* 1.3.2 (;^59^ RRID:SCR 016216), which is based on *Nipype* 1.1.9 (;^60^ RRID:SCR 002502).

#### Anatomical data preprocessing

The T1-weighted (T1w) image was corrected for intensity non-uniformity (INU) with N4Bias Field Correction,^61^ distributed with ANTs 2.2.0 (62, RRID:SCR 004757), and used as T1w-reference throughout the workflow. The T1w-reference was then skull-stripped with a *Nipype* implementation of the `antsBrainExtraction.sh` workflow (from ANTs), using OASIS30ANTs as target template. Spatial normalization to the *ICBM 152 Nonlinear Asymmetrical template version 2009c* was performed through nonlinear registration with `antsRegistration` (ANTs 2.2.0), using brain-extracted versions of both T1w volume and template. Brain tissue segmentation of cerebrospinal fluid (CSF), white-matter (WM) and gray-matter (GM) was performed on the brain-extracted T1w using `fast`.

#### Functional data preprocessing

For each of the 20 BOLD runs found per subject (across all tasks and sessions), the following preprocessing was performed. First, a reference volume and its skull-stripped version were generated using a custom methodology of *fMRIPrep*. A deformation field to correct for susceptibility distortions was estimated based on two echo-planar imaging (EPI) references with opposing phase-encoding directions, using `3dQwarp`(AFNI 20160207). Based on the estimated susceptibility distortion, an unwarped BOLD reference was calculated for a more accurate co-registration with the anatomical reference.

The BOLD reference was then co-registered to the T1w reference using `flirt` with the boundary-based registration cost-function. Co-registration was configured with nine degrees of freedom to account for distortions remaining in the BOLD reference. Head-motion parameters with respect to the BOLD reference (transformation matrices, and six corresponding rotation and translation parameters) are estimated before any spatiotemporal filtering using `mcflirt` The BOLD time-series (including slice-timing correction when applied) were resampled onto their original, native space by applying a single, composite transform to correct for head-motion and susceptibility distortions. These resampled BOLD time-series will be referred to as *preprocessed BOLD in original space*, or just *preprocessed BOLD*. The BOLD time-series were resampled to MNI152NLin2009cAsym standard space, generating a *preprocessed BOLD run in MNI152NLin2009cAsym space*. First, a reference volume and its skull-stripped version were generated using a custom methodology of *fMRIPrep*. Several confounding time-series were calculated based on the *preprocessed BOLD*: framewise displacement (FD), DVARS and three region-wise global signals. FD and DVARS are calculated for each functional run, both using their implementations in *Nipype*.

The three global signals are extracted within the CSF, the WM, and the whole-brain masks. Additionally, a set of physiological regressors were extracted to allow for component-based noise correction. Principal components are estimated after high-pass filtering the *preprocessed BOLD* time-series (using a discrete cosine filter with 128s cut-off) for the two *CompCor* variants: temporal (tCompCor) and anatomical (aCompCor). Six tCompCor components are then calculated from the top 5% variable voxels within a mask covering the subcortical regions. This subcortical mask is obtained by heavily eroding the brain mask, which ensures it does not include cortical GM regions. For aCompCor, six components are calculated within the intersection of the aforementioned mask and the union of CSF and WM masks calculated in T1w space, after their projection to the native space of each functional run (using the inverse BOLD-to-T1w transformation).

The head-motion estimates calculated in the correction step were also placed within the corresponding confounds file. All resamplings can be performed with *a single interpolation step* by composing all the pertinent transformations (i.e. head-motion transform matrices, susceptibility distortion correction when available, and co-registrations to anatomical and template spaces). Gridded (volumetric) resamplings were performed using ants Apply Transforms (ANTs), configured with Lanczos interpolation to minimize the smoothing effects of other kernels.

### Linear feature summation model (LFS model)

We hypothesized that subjective preferences for visual stimuli are constructed by the influence of visual and emotional features of the stimuli. As its simplest, we assumed that the subjective value of the *i*-th stimulus *v_i_* is computed by a weighted sum of feature values *f_i,j_*:

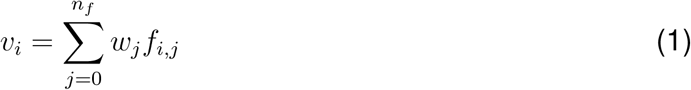

where *w_j_* is a weight of the *j*-th feature, *f_i,j_* is the value of the *j*-th feature for stimulus *i*, and *n_f_* is the number of features. The 0-th feature is a constant *f_i,_*_0_ = 1 for all *i*’s.

Importantly, *w_j_* is not a function of a particular stimulus but shared across all visual stimuli, reflecting the *taste* of a participant. The same taste (*w_j_*’s) can also be shared across different participants, as we showed in our behavioral analysis. The features *f_i,j_* were computed using visual stimuli; we used the same feature values to predict liking ratings across participants. We used the simple linear model Eq.(1) to predict liking ratings in our behavioral analysis (please see below for how we determined features and weights).

As we schematically showed in Figure 1, we hypothesized that the input stimulus was first broke down into low-level features and then transformed into high-level features, and indeed we found that a significant variance of high-level features can be predicted by a set of low-level features. This hierarchical structure of the LFS model was further tested in our DCNN and fMRI analysis.

### Features

Because we did not know apriori what features would best describe human aesthetic values for visual art, we constructed a large feature set using previously published methods from computer vision augmented with additional features that we ourselves identified using additional existing machine learning methods.

#### Visual low-level features introduced in^25^

We employed 40 visual features introduced in.^25^ We do not repeat descriptions of the features here; but briefly, the feature sets consist of 12 global features that are computed from the entire image that include color distributions, brightness effects, blurring effects, and edge detection, and 28 local features that are computed for separate segments of the image (the first, the second and the third largest segments). Most features are computed straightforwardly in either HSL (hue, saturation, lightness) or HSV (hue, saturation, value) space (e.g. average hue value).

One feature that deserves description is a blurring effect. Following,^25, 63^ we assumed that the image *I* was generated from a hypothetical sharp image with a Gaussian smoothing filter with an unknown variance *σ*. Assuming that the frequency distribution for the hypothetical image is approximately the same as the blurred, actual image, the parameter *σ* represents the degree to which the image was blurred. The *σ* was estimated by the Fourier transform of the original image by the highest frequency, whose power is greater than a certain threshold.

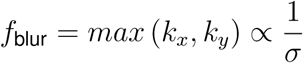

where *k_x_* = 2(*x − n_x_/*2)*/n_x_* and *k_y_* = 2(*y − n_y_/*2)*/n_y_* with (*x, y*) and (*n_x_, n_y_*) are the coordinates of the pixel and the total number of pixel values, respectively. The above max was taken within the components whose power is larger than four.^25^

The segmentation for this feature set was computed by a technique called kernel Graph-Cut.^28, 64^ Following,^25^ we generated a total of at least six segments for each image using a *C*^++^ and Matlab package for kernel graph cut segmentation.^64^ The regularization parameter that weighs the cost of cut against smoothness was adjusted for each image in order to obtain about six segments. Please see^25, 64^ for the full description of this method and examples.

Of these 40 features, we included all of them in our initial feature set except for local features for the third-largest segment, which were highly correlated with features for the first and second-largest segments and were thus deemed unlikely to add unique variance to the feature prediction stage.

#### Additional Low-Level Features

We assembled the following low-level features to supplement the set by Li & Chen.^25^ These include both global features and local features. Local features were calculated on segments determined by two methods. The first method was statistical region merging (SRM) as implemented by,^65^ where the segmentation parameter was incremented until at least three segments were calculated. The second method converted paintings into LAB color space and used k-means clustering of the A and B components. While the first method reliably identified distinct shapes in the paintings, the second method reliably identified distinct color motifs in the paintings. Examples of the two segmentation methods are given in Figure S14.

The segmentation method for each feature is indicated in the following descriptions. Each local feature was calculated on the first and second-largest segments.

Local Features:

• Segment Size (SRM): Segment size for segment *i* was calculated as the area of segment i over the area of the entire image:

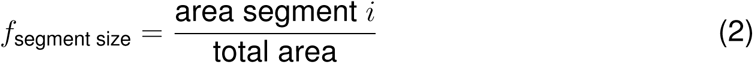

• HSV Mean (SRM): To calculate mean hue, saturation and color value for each segment, segments were converted from RGB to HSV color space.

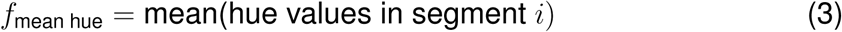

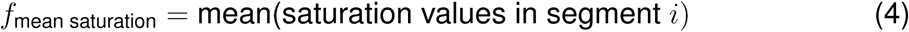

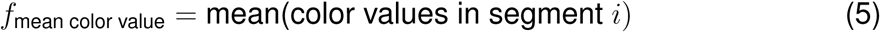

• Segment Moments (SRM):

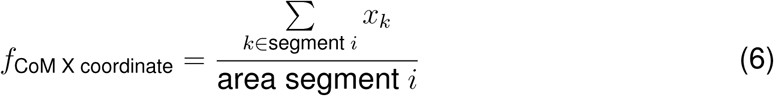

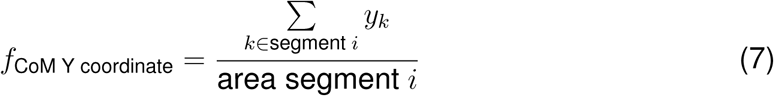

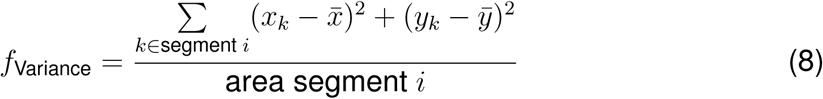

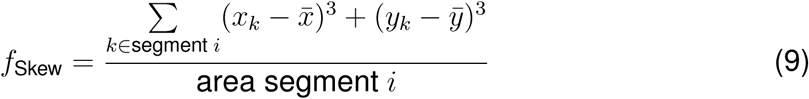

where 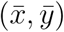 is the center of mass coordinates of the corresponding segment.

• Entropy (SRM):

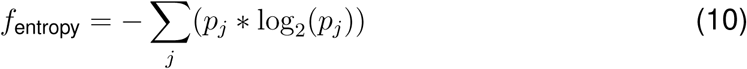

where *p* equals the normalized intensity histogram counts of segment *i*.

• Symmetry (SRM): For each segment, the painting was cropped to maximum dimensions of the segment. The horizontal and vertical mirror images of the rectangle were taken, and the mean squared error of each was calculated from the original.

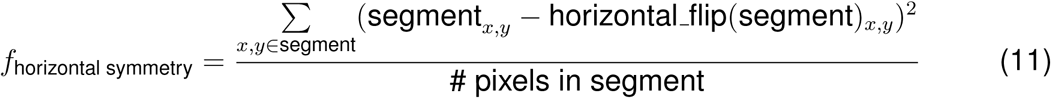

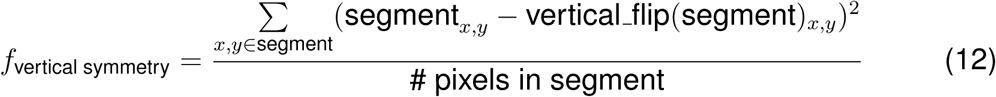

• R-Value Mean (K-Means): Originally, we took the mean of R, G, and B values for each segment, but found these values to be highly correlated, so we reduced these three features down to just one feature for mean R value.

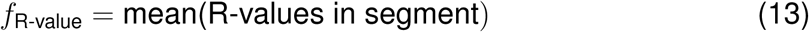

• HSV Mean (K-Means): As with SRM generated segments, we took the hue, saturation, and color value means of segments generated by K-means segmentation as described in equations 2-4.

Global Features:

• Image Intensity: Paintings were converted from RGB to grayscale from 0 to 255 to yield a measure of intensity. The 0-255 scale was divided into five equally-sized bins. Each bin count accounted for one feature.

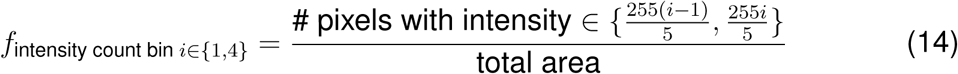

• HSV Modes: Paintings were converted to HSV space, and the modes of the hue, saturation, and color value across the entire painting were calculated. While we took mean HSV values over segments in an effort to calculate overall-segment statistics, we took the mode HSV values across the entire image in an effort to extract dominating trends across the painting as a whole.

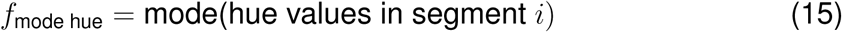

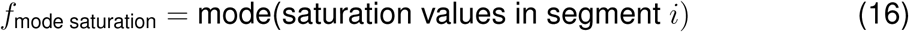

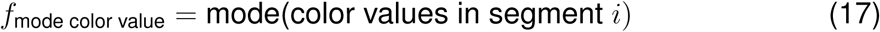

• Aspect (width-height) Ratio:

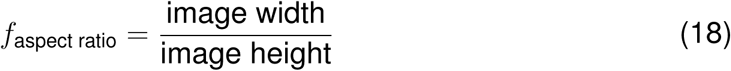

• Entropy: Entropy over the entire painting was calculated according to equation 9.

In addition, we also annotated our image set with whether or not each image included a person. This was done by manual annotation, but it can also be done with a human detection algorithm (e.g., see^66^). We included this presence-of-a-person feature in the low-level feature set.

#### High-Level Feature Set^26, 27^

We also introduced features that are more abstract and not easily computed by a simple algorithm. In,^26^ Chatterjee et al. pioneered this by introducing 12 features (color temperature, depth, abstract, realism, balance, accuracy, stroke, animacy, emotion, color saturation, complexity) that were annotated by human participants for 24 paintings, in which the authors have found that annotations were consistent across participants, regardless of their artistic experience. Vaidya et al.^27^ further collected annotations of these feature sets from artistically experienced participants for an additional 175 paintings and performed a principal component analysis, finding three major components that summarize the variance of the original 12 features. Inspired by the three principal components, we introduced three high-level features: concreteness, dynamics, and temperature. Also, we introduced valence as an additional high-level feature ^1^. The four high-level features were annotated in a similar manner to the previous studies.^26, 27^We took the mean annotations of all 13 participants for each image as feature values.

### Identifying the shared feature set that predicts aesthetic preferences

The above method allowed us to have a set of 83 features in total that are possibly used to predict human aesthetic valuation. These features are likely redundant because some of them are highly correlated, and many may not contribute to decisions at all. We thus sought to identify a minimal subset of features that are commonly used by participants. We performed this analysis using Matlab Sparse Gradient Descent Library ^2^. For this, we first orthogonalized features by sparse PCA.^67^ Then we performed a regression with a LASSO penalty at the group level using the seven in-lab participants’ behavioral data with a function *group lasso problem*. We used Fast Iterative Soft Thresholding Algorithm (FISTA) with cross-validation. After eliminating PC’s that were not shared by more than one participant, we transformed the PC’s back to the original space. We then eliminated one of the two features that were most highly correlated (*r*^2^ *>* 0.5) to obtain the final set of shared features.

The identified shared features from the in-lab behavioral participants are the following: the concreteness, the dynamics, the temperature, the valence, the global hue contrast from,^25^ the global brightness contrast from,^25^ the blurring effect from,^25^ the vertical center of largest segment using the Graph-cut from,^25^ the average saturation of the second-largest segment using the Graph-cut from,^25^ the blurring contrast between the largest and the second largest segments using the Graph-cut in,^25^ the size of largest segment using SRM, width-height ratio, and the presence of a person.

For the fMRI analysis, we used behavioral data from both in-lab and fMRI participants (13 particiapnts in total). Because the goal of the fMRI analysis is to highlight the hierarchical nature in neural coding between low and high-level features, we first repeated the above procedure with low-level features alone (79 features in total) and then we added high-level features (the concreteness, the dynamics, the temperature, and the valance) to the obtained shared low-level features.

The identified shared features for the fMRI analysis using behavioral data from the in-lab and fMRI particiapnts are the following: the concreteness, the dynamics, the temperature, the valence, the global average saturation from,^25^ the global blurring effect from,^25^the horizontal coordinate of mass center for the largest segment using the Graph-cut from,^25^ the vertical coordinate of mass center for the largest segment using the Graph-cut from,^25^ the mass skewness for the second largest segment using the Graph-cut from,^25^ the size of the largest segment using SRM, the mean hue of the largest segment using SRM, the mean color value of largest segment using SRM, the mass variance of the largest segment using SRM, global entropy, the entropy of the second-largest segment using SRM, the image intensity in bin 1, the image intensity in bin 2, and the presence of a person.

### Representation similarity analyses

We computed a representation dissimilarity matrix over visual art stimuli using low-level or high-level features alone. The distance was measured by one minus correlation.

### Model fitting

We tested how our shared-feature model can predict human liking ratings using out-of-sample tests. All models were cross-validated in twenty folds, and we used ridge regression unless otherwise stated. Hyperparameters were tuned by cross-validation. In one analysis, we trained our regression model with a shared feature set on the average ratings of six in-lab participants and tested on the remaining participant. We calculated the Pearson correlation between model predictions (pooled predictions from all cross-validation sets) and actual data, and defined it as the predictive accuracy. In another analysis, we trained our regression model on one participant and tested on the same participant (randomly partitioned to 20 folds). In a further analysis, we also trained our model on the average ratings of on-line participants and tested it on in-lab participants, thereby ensuring complete independent between our training data and testing data. We also tested our model on on-line participants. In one analysis, we trained our model on *n* − 1 participants and tested on the remaining participant (leave-one-out). In another, we trained our model on in-lab participants and tested on-line participants. We also tested our model with low-level features only. For this, we trained our model with all low-level features with a lasso penalty on *n* − 1 on-line participants and tested on the remaining online participant.

We also estimated individual participant’s feature weights by fitting a linear regression model with the shared feature set to each participant. For illustrative purposes, the weights were normalized for each participant by the maximum feature value (concreteness) in Figures 1G, S3, S4.

We further tested whether the low-level feature set could predict high-level features. For this, we used a support vector machine with cross-validation (fitcsvm in Matlab).

We performed the same analysis with our dataset for photo contest images (AVA). Because we do not have high-level feature ratings fro the photo contest database and we also did not obtain manual ratings of the presence of a person for that image database, we only used low-level features (minus the presence of a person feature) for analyzing this data. In one analysis, we trained our model on the AVA dataset and tested it on the AVA dataset. In another analysis We trained our model on the average ratings of online ART dataset and tested it on the AVA dataset. Also, we trained our model on the AVA dataset ratings and tested it on the online ART dataset ratings. We used all low-level features with a strong lasso penalty and cross-validation to avoid over-fitting. The hyperparameters were optimized by cross-validation.

The significance of the above analyses was measured by generating a null distribution constructed by the same analyses but with permuted image labels. The null distribution was construed by 10000 permutations. The chance level was determined by the mean of the null distribution.

### Cluster analysis

We explored individual differences in the feature space by applying a clustering analysis of on-line participants. For this, we first transformed our shared-feature set to principal component (PC) space and fit our model in the PC space. The obtained weights were normalized by using each participant’s weight with the maximum magnitude. Then we fit a Gaussian mixture model with a different number of nodes to the resulting weights, assuming that the off-diagonal terms of the covariance are zero because the fit is in the PC space. We used an expectation-maximization method with 100 different random starting points, using Matlab’s function fitgmdist. We compared the Bayesian Information Criterion scores for each fit. We then compared the results of models with the number of clusters set to n=1,2,3,4,5,6 and took the model with the minimum BIC score. This turned out to be a model with n=3 clusters. Hard clustering was used, where each data point in the weight space was assigned to the component yielding the highest posterior probability.

### Deep Convolutional Neural Network (DCNN) analysis

#### Network architecture

The deep convolutional neural network (DCNN) we used consists of two parts. An input image feeds into convolutional layers from the standard VGG-16 network that is pre-trained on ImageNet. The output of the convolutional layers then projects to fully connected layers. This architecture follows the current state-of-the-art model on aesthetic evaluation.^29, 68^

The details of the convolutional layers from the VGG network can be found in;^30^ but briefly, it consists of 13 convolutional layers and 5 intervening max pooling layers. Each convolutional layer is followed by a rectified linear unit (ReLU). The output of the final convolutional layer is flattened to a 25088-dimensional vector so that it can be fed into the fully connected layer.

The fully connected part has two hidden layers, where each layer has 4096 dimensions. The fully connected layers are also followed by a ReLU layer. During training, a dropout layer was added with a drop out probability 0.5 after every ReLU layer for regularization. Following the current state of the art model,^68^ the output of the fully connected network is a 10-dimensional vector that is normalized by a softmax. The output vector was weighted averaged to produce a scalar value^68^ that ranges from 0 to 3.

#### Network training

We trained our model on our in-lab behavioral data set by tuning weights in the fully connected layers. Training all layers on the ART and/or AVA data set are also possible, but we found that the model’s predictive performance does not improve even if we trained all layers. We employed 10-fold cross-validation to benchmark the art rating prediction.

The model was optimized using a Huber loss metric, which is robust to outliers.^69^ The average rating of each image among the in-lab participants was used as the ground truth.

We used stochastic gradient descent (SGD) with momentum to train the model. We used a batch size of 100, a learning rate of 10*^−^*^4^, the momentum of 0.9, and weight decay of 5 × 10*^−^*^4^. The learning rate decayed by a factor of 0.1 every 30 epochs.

To handle various sizes of images, we used the zero-padding method. Because our model could only have a 224 × 224 sized input, we first scaled the input images to have the longer edges be 224 pixels long. Then we filled the remaining space with 0 valued pixels (black).

We used Python 3.7, Pytorch 0.4.1.post2, and CUDA 9.0 throughout the analysis.

#### Retraining DCNN to extract hidden layer activations

We also trained our network on a single fold ART data in order to obtain a single set of hidden layer activations. We prevented over-fitting by stopping our training when the model performance (Pearson correlation between the model’s prediction and data) reached the mean correlation from the 10-folds cross-validation.

#### Decoding features from the deep neural network

We decoded the LFS model’s features from hidden layers by using linear (for continuous features) and logistic (for categorical features) regression models. We considered the activations of outputs of ReLU layers (total of 15 layers). First, we split the data into ten folds for the 10-fold cross-validation. In each iteration of the cross-validation, because dimensions of the hidden layers are much larger (64 × 224 × 224 = 3211264) than the actual data size, we first performed PCA on the activation of each hidden layer from the training set. The number of principal components was chosen to account for 80% of the total variance. By doing so, each layer’s dimension was reduced to less than 536. Then the hidden layers’ activations from the test set were projected onto the principal component space by using the fitted PCA transformation matrices. The regularization coefficient for the regression was tuned by doing a grid search, and the best performing coefficient for each layer and feature was chosen based on the scores from the 10-folds cross-validation. We tested for a total of 19 features, including all 18 features that we used for our fMRI analysis, as well as the simplest feature that was not included into our fMRI analysis (as a result of our group-level feature selection) but that was also of interest here: the average hue value. In a supplementary analysis, we also explored whether adding ‘style matrices’ of hidden layers^70^ to the PCA-transformed hidden layer’s activations can improve the decoding accuracy; however, we found the style matrices do not improve the decoding accuracy. Sklearn 0.19.2 on Python 3.7 was used.

#### Reclassifying features according to the slopes of the doodling accuracy across hidden layers

In our LFS model, we classified putative low-level and high-level features simply by whether a feature is computed by a computer algorithm vs annotated by humans respectively. In reality, however, some putative low-level features are more complex in terms of how they could be constructed than other lower level features, while some putative high-level features could in fact be computed straightforwardly from raw pixel inputs. Using the decoding results of the features from hidden layers in the DCNN, we identified DCNN-defined low-level and high-level features. For this, we fit a linear slope to the estimated decoding accuracy vs hidden layers. We permuted layer labels 10,000 times and performed the same analysis to construct null distribution as described earlier. We classified a feature as high-level if the slope was significantly positive at *p <* 0.001, and we classified a feature as a low-level feature if the slope was signifcantly negative at *p <* 0.001.

The features showing negative slopes were: the average hue, the average saturation, the average hue of the largest segment using GraphCut, the average color value of the largest segment using GraphCut, the image intensity in bin 1, the image intensity in bin 3, and the temperature.

The features showing positive slopes were: the concreteness, the dynamics, the presence of a person, the vertical coordinate of the mass center for the largest segment using the Graph Cut, the mass variance of the largest segment using the SRM, the entropy in the 2nd largest segment using SRM. All of these require relatively complex computations, such as localization of segments or image identification. This is consistent with a previous study showing that object-related local features showed a similar increased decodability at a deeper layer.^32^

### fMRI analysis

We conducted fMRI data analysis with SPM 12. SPM’s feature for asymmetrically orthogonalizing parametric regressors was disabled throughout our analysis. We collected enough data from each individual participant (four days of scanning) so that we can analyze and interpret each participant’s results separately. The following regressors were obtained from the fmriprep preprocessing pipeline added to all analysis as nuisance regressors: frame-wise displacement, comp-cor, non-steady, trans, rot. The onsets of Stimulus, Decision, and Action were also controlled by stick regressors in all GLMs described below. In addition, we added the onset of the Decision period, the onset of feedback to all GLM as nuisance regressors, because we focused on the stimulus presentation period.

#### Identifying subjective value coding (GLM 1)

In order to gain insight into how the subjective value of art was represented in the brain, we performed a simple GLM analysis with a parametric regressor at the onset of Stimulus (GLM 1). The parameter was linearly modulated by participant’s liking ratings on each trial. The results were cluster FWE collected with a height threshold of *p <* 0.001.

#### Identifying feature coding (GLM 2,3, 2’, 3’)

In order to gain insight into how features were represented in the brain, we performed another GLM analysis with a parametric regressor at the onset of Stimulus (GLM 2,3). In GLM 2, there are in total 18 feature-modulated regressors; each representing the value of one of the shared features for the fMRI analysis. We then performed F-tests on high-level features and low-level features (a diagonal contrast matrix with an entry set to one for each feature of interest was constructed in SPM) in order to test whether a voxel is significantly modulated by any of the high and/or low-level features. We then counted the number of voxels that are significantly correlated (*p <* 0.001) in each ROI (note that the F-value for significance is different for high and low features due to the difference in the number of consisting features). We then displayed the proportions of two numbers in a given ROI.

We performed a similar analysis using the DCNN’s hidden layers (GLM 3). We took the first three principal components of each convolutional and fully connected layers (three PCs times 15 layers = 45 parametric regressors). We then performed F-tests on PCs from layers 1 to 4, layers 5 to 9, layers 10 to 13, and fully connected layers (layers 14 and 15). The proportions of the survived voxels were computed for each ROI.

In addition, we also performed the same analyses with GLMs to which we added liking ratings for each stimulus. We call these analyses GLM 2’ and GLM 3’, respectively.

#### Region of Interests (ROI)

We constructed ROIs for visual topographic areas using a previously published probabilistic map.^41^ We constructed 17 masks based on the 17 probabilistic maps taken from,^41^ consisting of 8 ventral–temporal (V1v, V2v, V3v, hV4, VO1, VO2, PHC1, and PHC2) and 9 dorsal–lateral (V1d, V2d, V3d, V3A, V3B, LO1, LO2, hMT, and MST) masks. In this, ventral and dorsal regions for early visual areas V1, V2, V3 are separately defined. Each mask was constructed by thresholding the probability map at *p >* 0.01. We defined *V*_123_ as V1v +V2v+ V3v+ V1d + V2d + V3d + V3A + V3B, and *V_high_* as hV4 + VO1 + VO2 + PHC1 + PHC2 + LO1 + LO2 + hMT + MST. (hV4: human V4, VO: ventral occipital cortex, PHC: posterior parahippocampal cortex, LO: lateral occipital cortex, hMT: human middle temporal area, MST: medial superior temporal area.)

We also constructed ROIs for parietal and prefrontal cortices using the AAL database. Posterior parietal cortex (PPC) was defined by bilateral MNI-Parietal-Inf + MNI-Parietal-Sup. lateral rbitofrontal cortex (lOFC) was defined by bilateral MNI-Frontal-Mid-Orb + MNI-Frontal-Inf-Orb + MNI-Frontal-Sup-Orb, and medial OFC (mOFC) was defined by bilateral MNI-Frontal-Med-Orb + bilateral MNI-Rectus. Dorsomedial PFC (dmPFC) was defined by bilateral MNI-Frontal-Sup-Medial + MNI-Cingulum-Ant, and dorsolateral PFC (dlPFC) was defined by bilateral MNI-Frontal-Mid + MNI-Frontal-Sup. Ventrolateral PFC (vlPFC) was defined by bilateral MNI-Frontal-Inf-Oper + MNI-Frontal-Inf-Tri.

We also constructed lateral PFC (LPFC) as vlPFC + dlPFC +lOFC, and medial PFC (MPFC) as mOFC + dmPFC.

#### PPI analysis (GLM 4, 4’)

We conducted a psychobiological-physiological interaction analysis. We took a seed from GLM1’s cluster showing subjective value in MPFC (Figure S10), and psychological regressor as a box function, which is set to one during the stimulus epoch and 0 otherwise. We added the time course of the seed, the PPI regressor, to a variant of GLM 2 (the parametric regressors in which feature values were constructed using a boxcar function at stimulus periods, instead of its onsets) and determined which voxels were correlated with the PPI regressor (GLM 4). Following,^17^ boxcar functions were used because feature integration can take place throughout the duration of each stimulus presentation.

We also conducted a control PPI analysis. For this we took the same seed, but now the psychological regressor was a box function, which is one during ITI and 0 otherwise. We added the time course of the seed and the PPI regressor, the box function for ITI, and the PPI regressor to the same variant of GLM 2 (the parametric regressors with feature values were constructed using boxcar function at Stimulus periods, instead of its onsets). We refer to this as GLM 4’.

#### feature integration analysis

We conducted an F-test using GLM 2, asking any of the shared features was significantly correlated with a given voxel (a diagonal with one at all features in SPM). The resulting F-map is thresholded at *p <* 0.05 cFWE at the whole-brain with height threshold at *p <* 0.001. We then asked within the survived voxels, which of them were also significantly positively correlated with PPI regressor in GLM 4, using at value thresholded at *p <* 0.001 uncorrected. We then counted the fraction of voxels that survived this test in a given ROI.

## Acknowledgements

We thank Peter Dayan, Shin Shimojo, Pietro Perona, Lesley Fellows, Avinash Vaidya, Jeff Cockburn and Logan Cross for discussions and suggestions. This work was supported by NIDA grant R01DA040011 and the Caltech Conte Center for Social Decision Making (P50MH094258) to JOD, the Japan Society for Promotion of Science the Swartz Foundation and the Suntory Foundation to KI, and the William H. and Helen Lang SURF Fellowship to IW.

## Author Contributions

K.I. and J.P.O. conceived and designed the project. K.I., S.Y., I.A.W., K.T., performed experiments and K.I., S.Y., I.A.W., K.T., J.P.O. analyzed and discussed results. K.I., S.Y., I.A.W., J.P.O. wrote the manuscript.

## Supplementary figures

**Figure S1:**
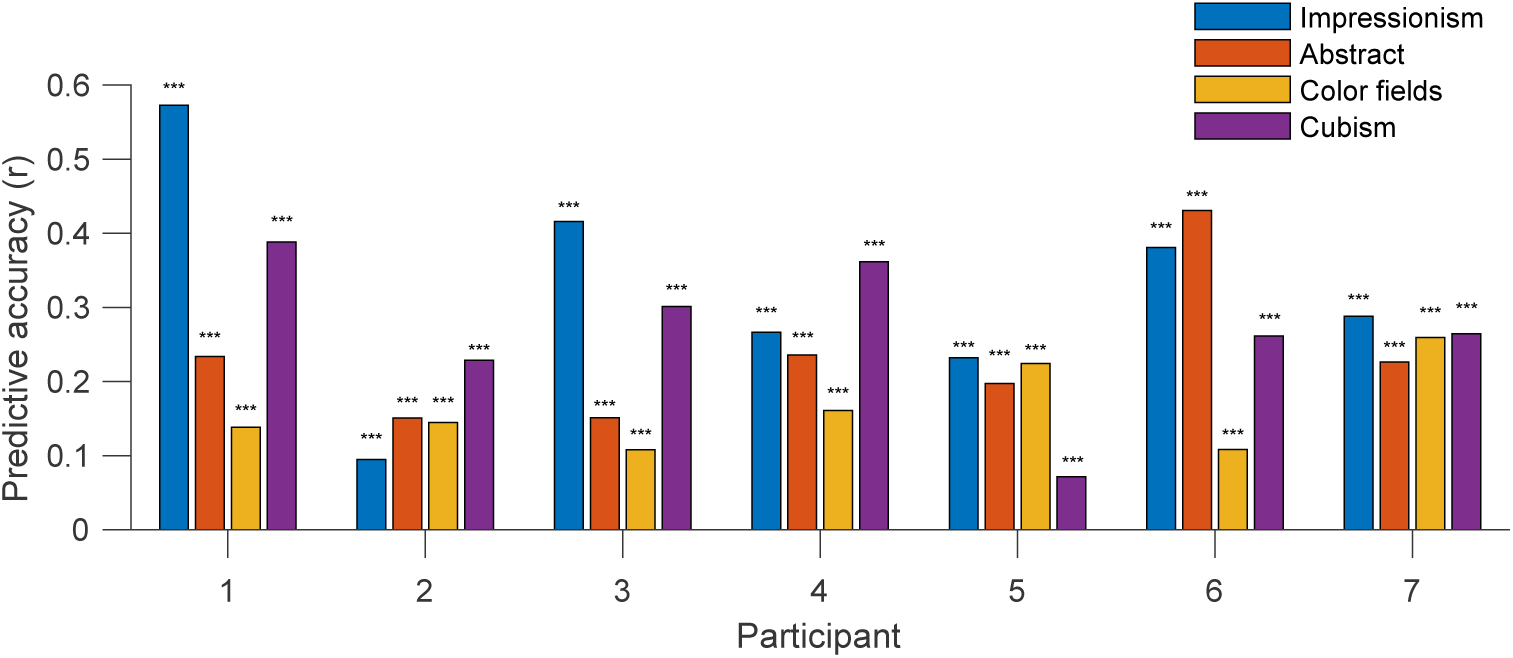
Predictive accuracy of the LFS model within different art genres. The model was trained on all images using 20 fold cross validation in each participant. Predictions for images in each art genre were compared with the actual data. The predictive accuracy was measured by Pearson correlation. This figure shows that our overall predictive accuracy is not merely an artifact of the fact that people like different genres differently, i.e. that the LFS model is sensitive only to differences between images as a result of genre and that this alone enables it to have success. Here, even within specific genres, the model can still succeed in predicting liking ratings just as it can across genres. Note that correlation values are smaller than the overall value presented in Figure 1. This is because between-genre correlation is indeed present in Figure 1.

**Figure S2:**
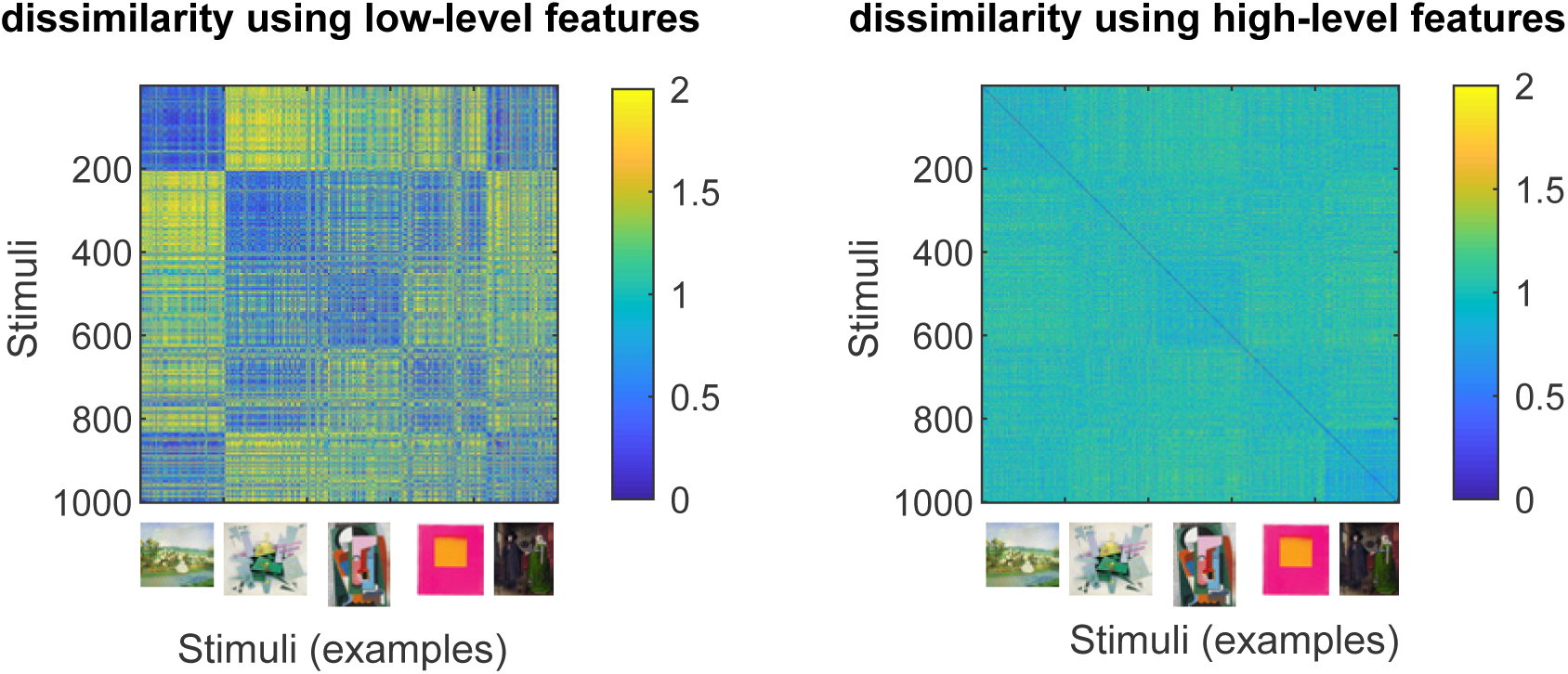
Representation dissimilarity matrix using low-level and high-level features. Image index; 1-204: Impressionism. 205-417: Abstract art. 418-621: Color fields. 622-826: Cubism. 827-1000: Pictures from the stimulus set of.^27^ Example stimuli from different art genres are shown in x axis.

**Figure S3:**
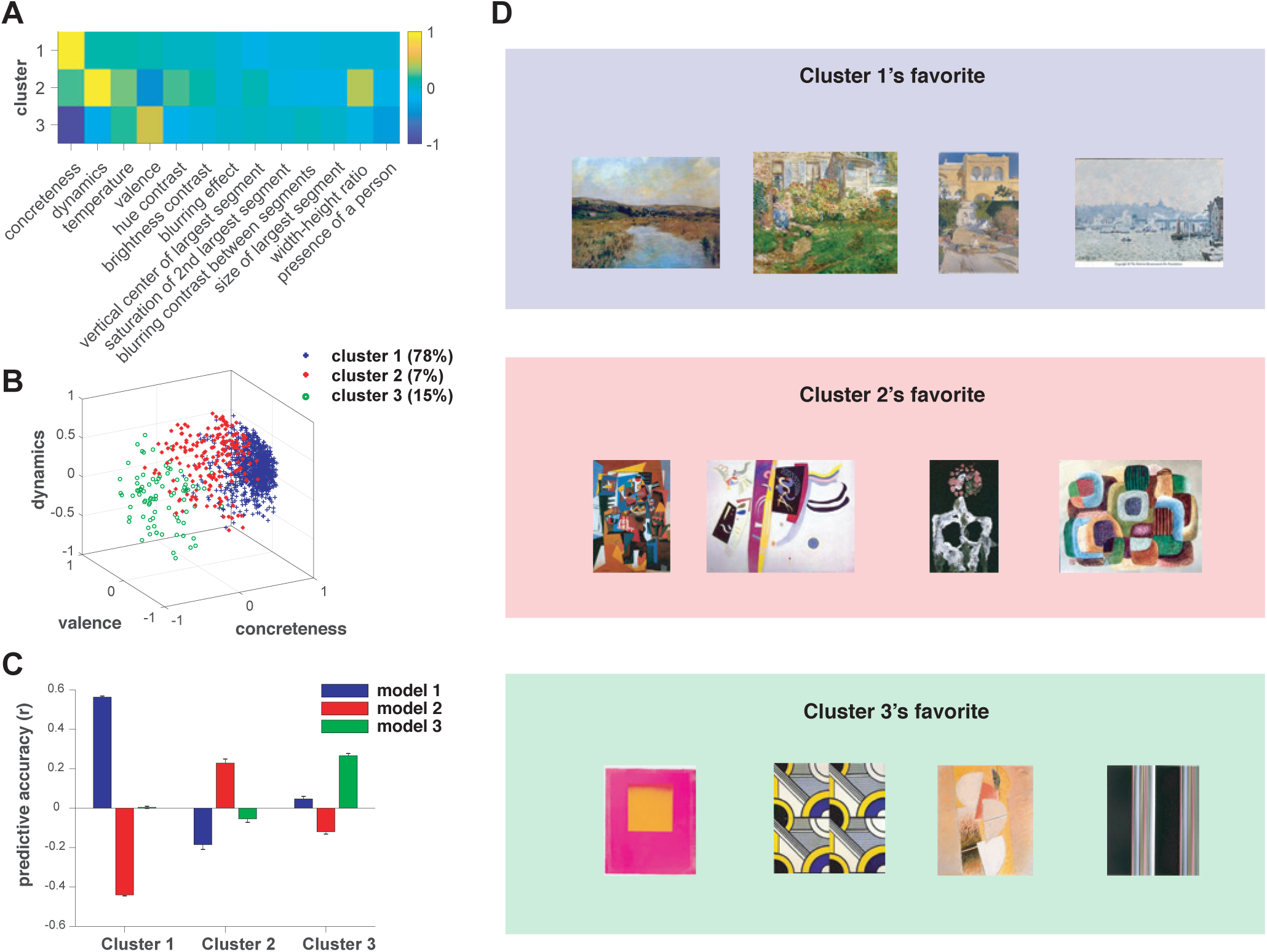
Clustering analysis. We fit the LFS model to each individual online participant. We then performed a clustering analysis on the estimated weights using a Gaussian mixture model. The number of Gaussians was optimized by comparing Bayes Information Criterion (BIC) scores. (**A**). The estimated feature weights at the center of each cluster. (**B**). The estimated feature weights of all participants, colored by cluster membership. (**C**). Predictive accuracy of participants in each cluster, using the model with the means of each gaussian as its parameters. (**D**). Example stimuli that were preferred by each cluster.

**Figure S4:**
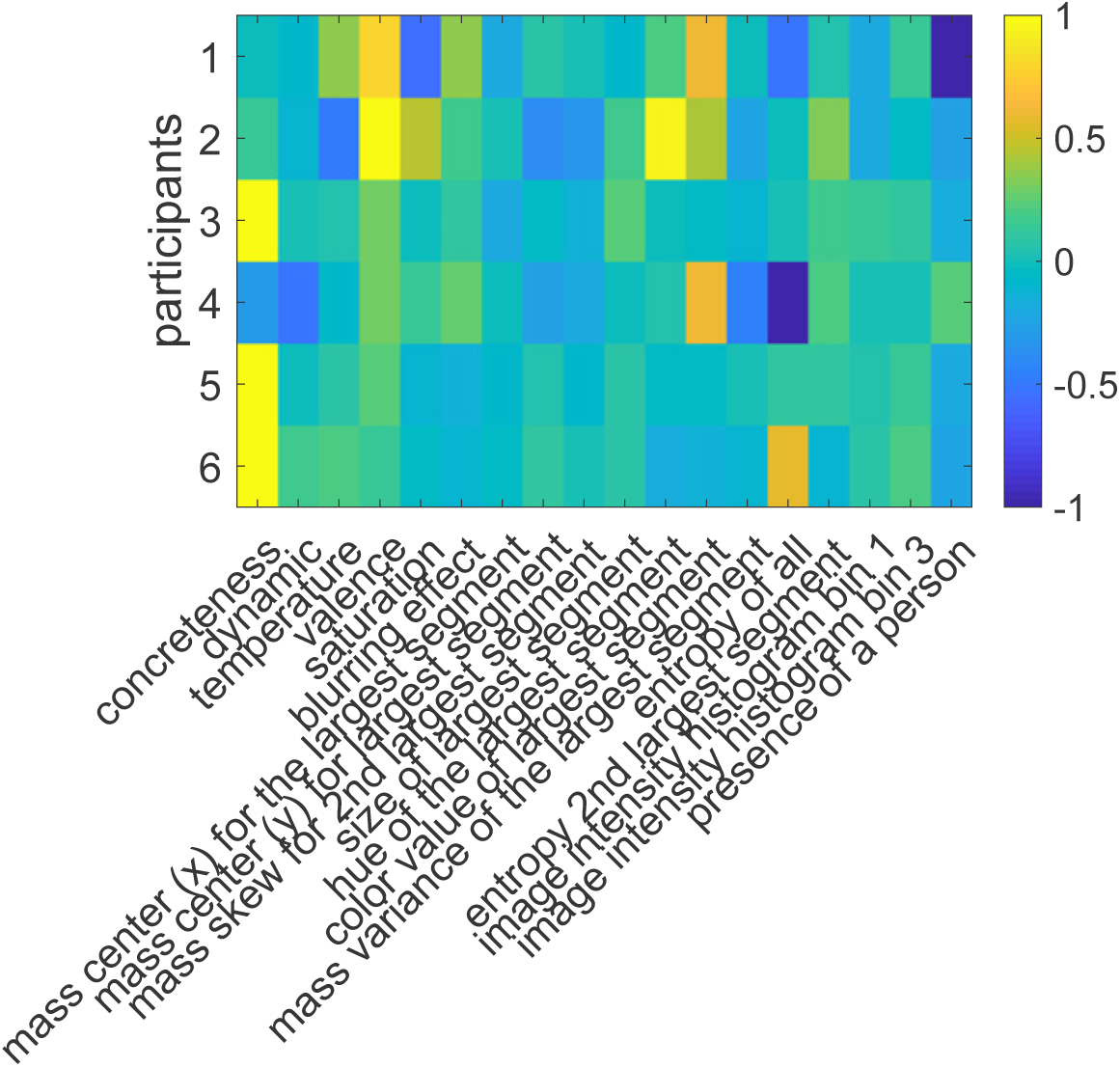
Feature weights from each fMRI participant, determined by fitting the LFS model to the liking ratings from each participant.

**Figure S5:**
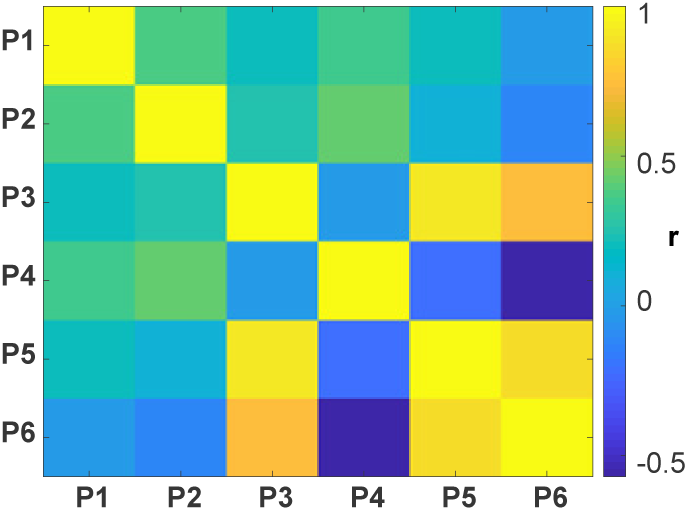
Correlations in feature weights between fMRI participants.

**Figure S6:**
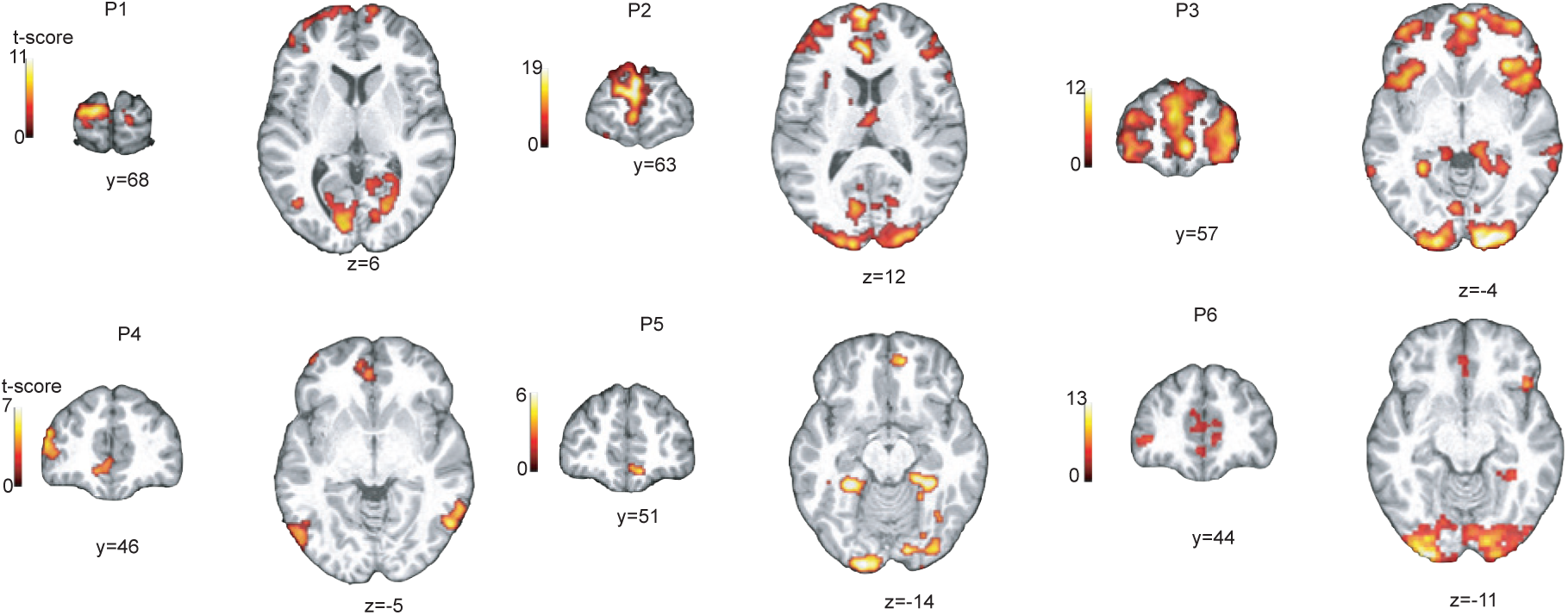
Neural correlates of subjective value. Clusters at whole-brain cFWE *p <* 0.05 with height threshold at *p <* 0.001 are shown.

**Figure S7:**
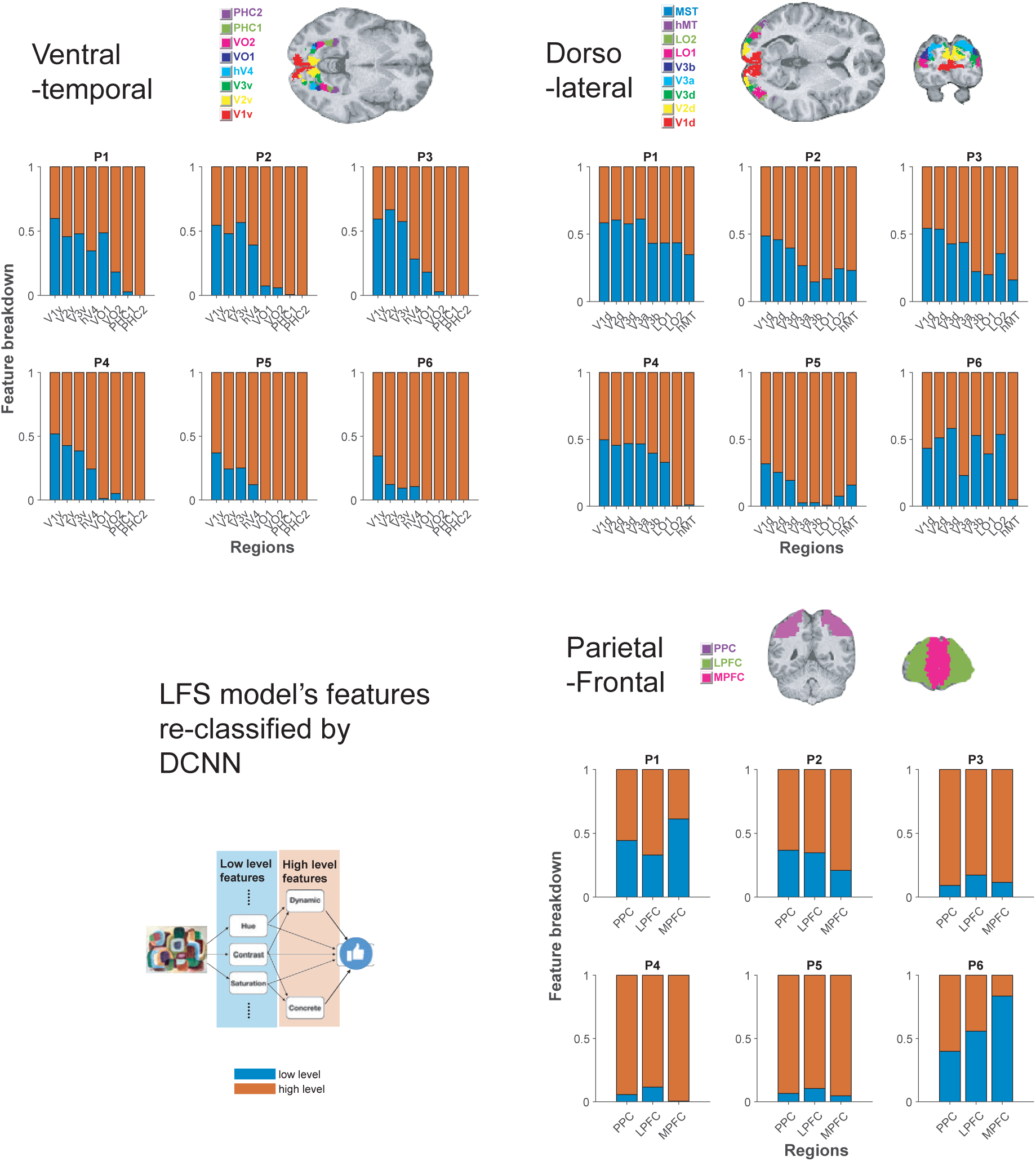
Results of fMRI encoding analysis of low- and high-level features, using the features that are reclassified according to the DCNN results. Among the features that were originally considered, the features showing significantly positive slopes across layers in the DCNN were defined as high-level features, while the features showing significantly negative slopes across layers in the DCNN were defined as low-level features. The results did not qualitatively change from our original analysis with the original definition of low- and high-level features.

**Figure S8:**
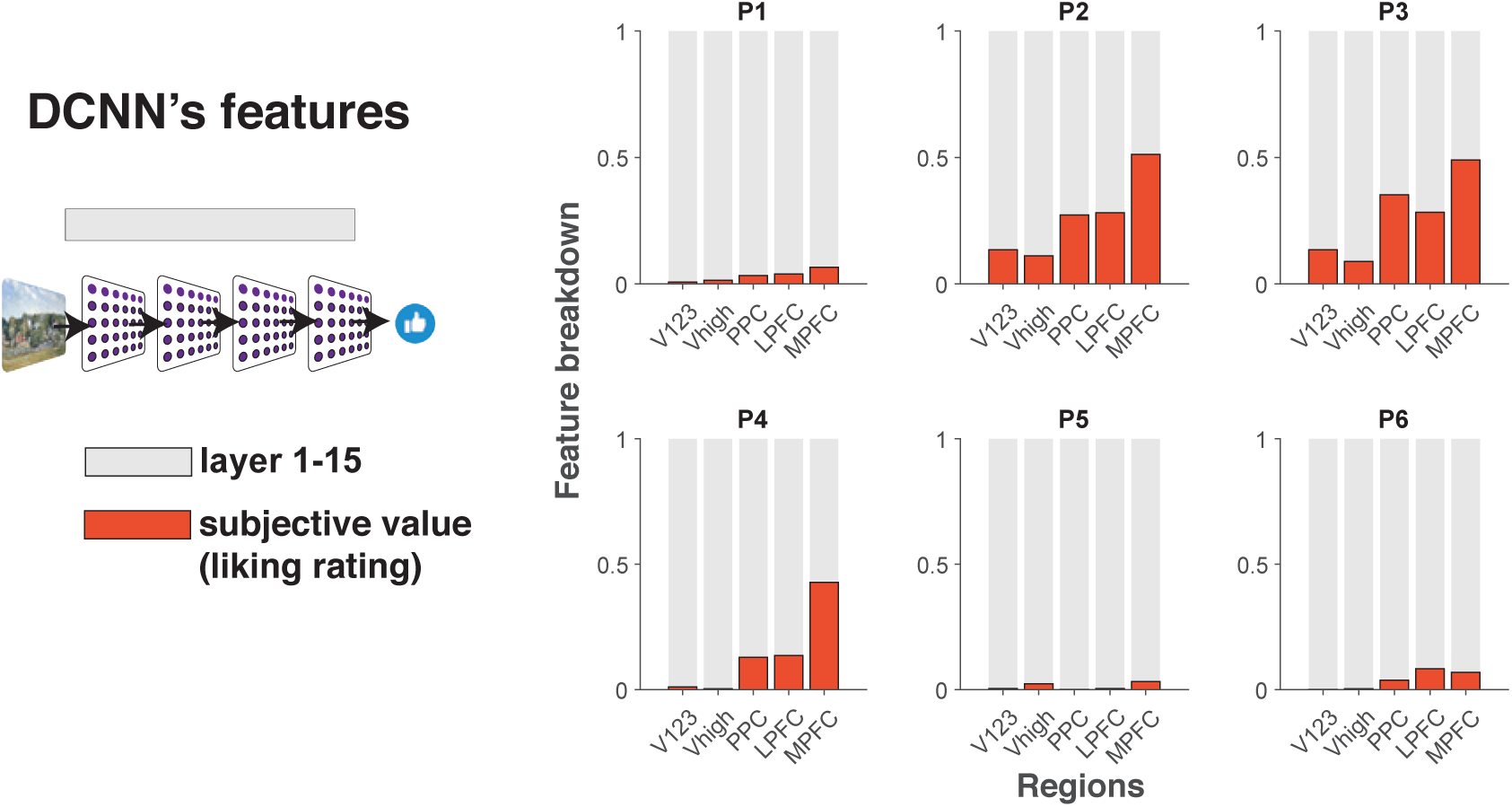
The same analysis as Figure 7A but now with DCNN model’s features.

**Figure S9:**
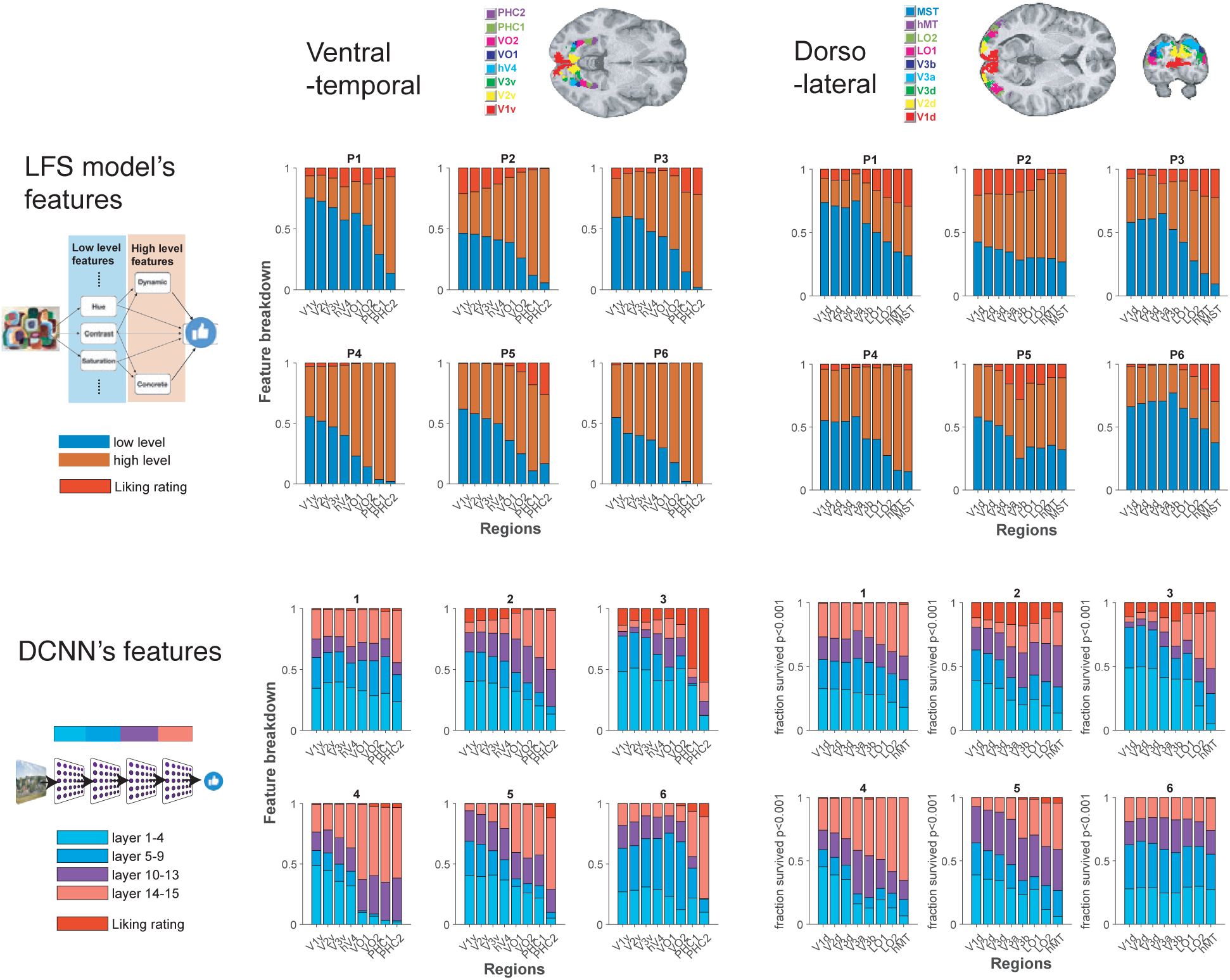
The encoding analysis of low- and high-level features when subjective liking ratings were also included into the same GLM. The results of ROIs in the ventral-temporal and dorso-lateral visual streams are shown.

**Figure S10:**
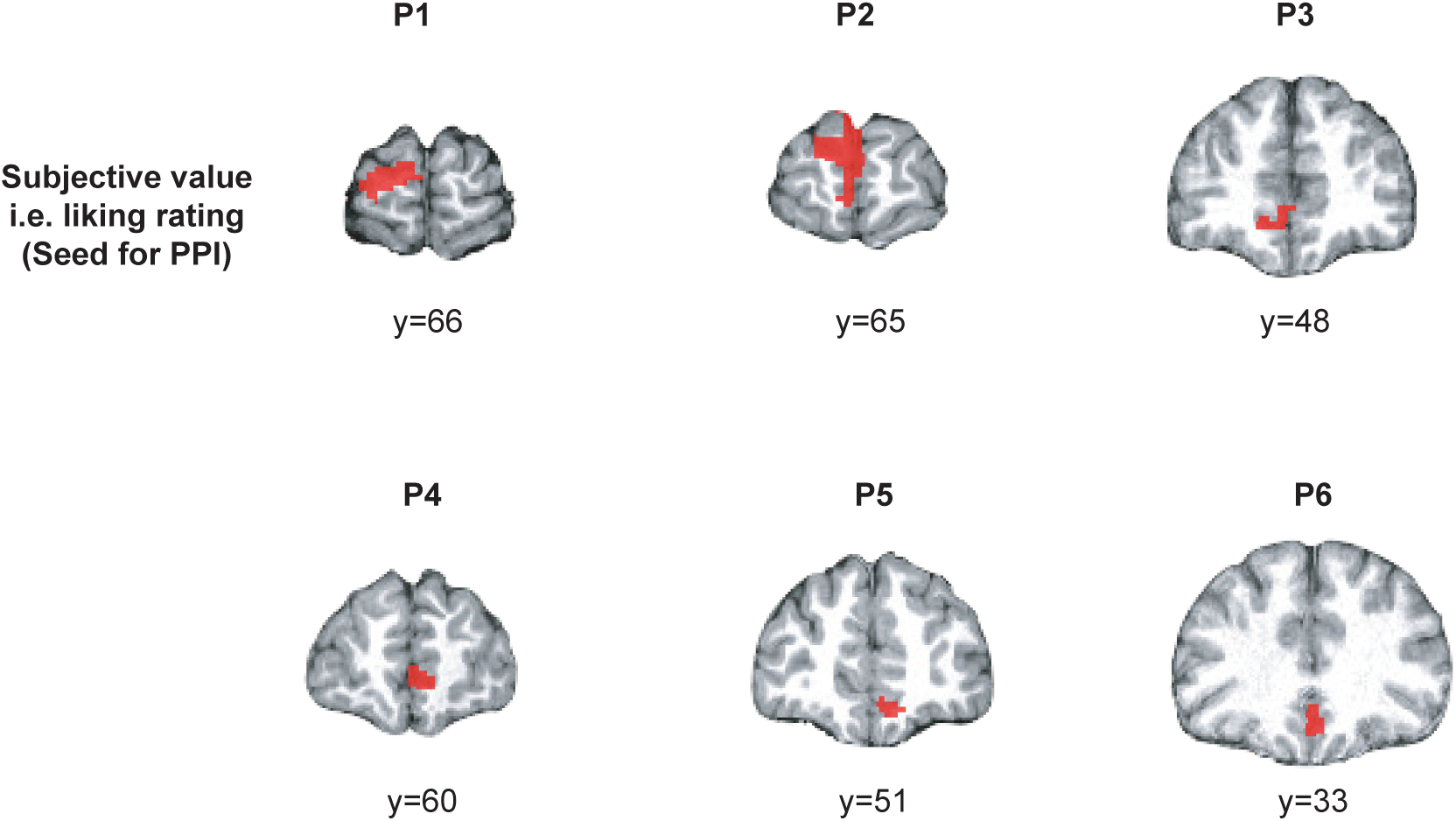
The seeds of the PPI analysis. The seeds used for the PPI analysis corresponded to a medial PFC cluster drawn from each participant that was found to show significant correlation with subjective value. The cluster used for each participant is shown here (*p <* 0.05 cFWE at whole-brain with a height threshold of *p <* 0.001).

**Figure S11:**
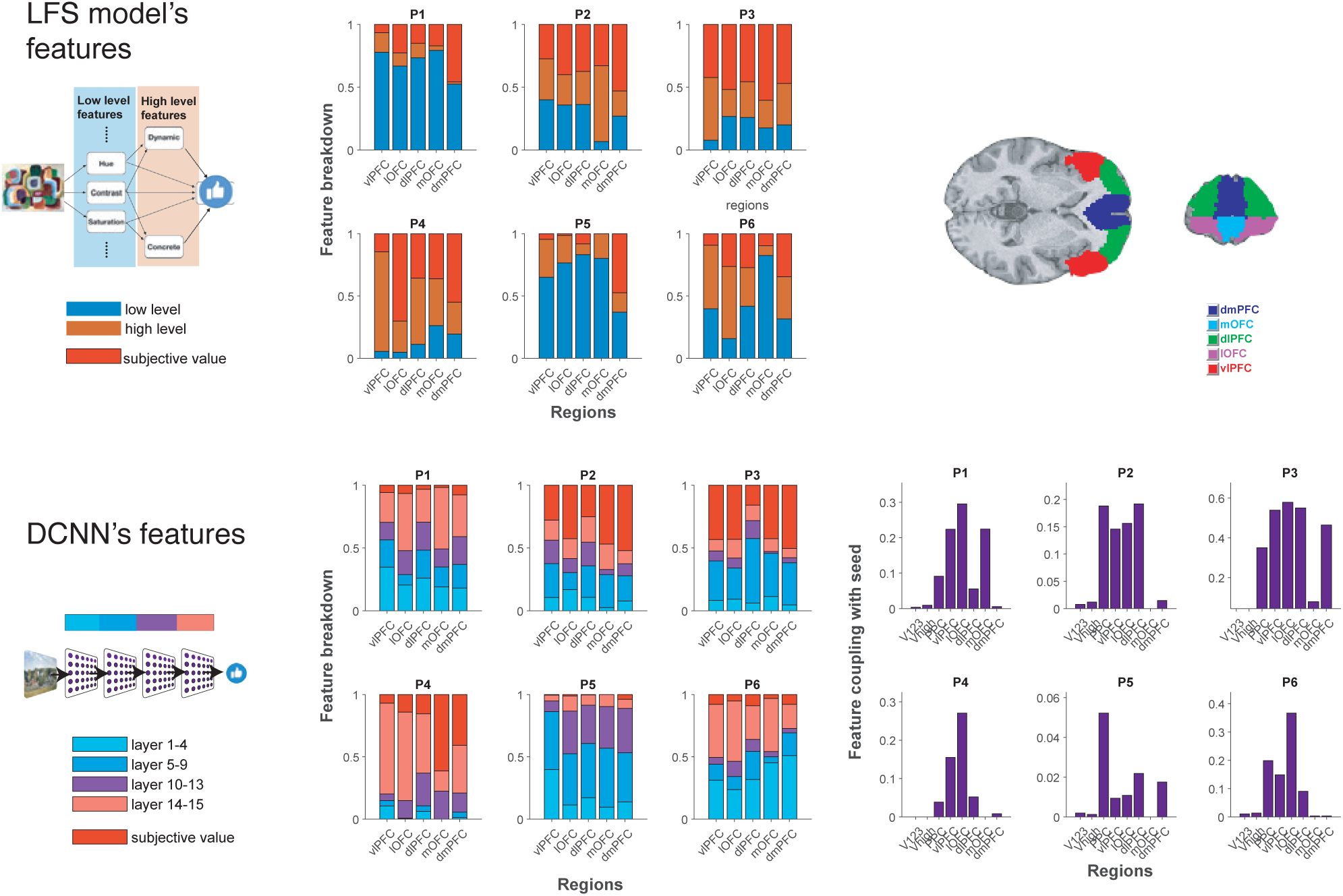
The same results as in Figure 7 and S8, but now broken down to show separate results for each sub-region of the PFC.

**Figure S12:**
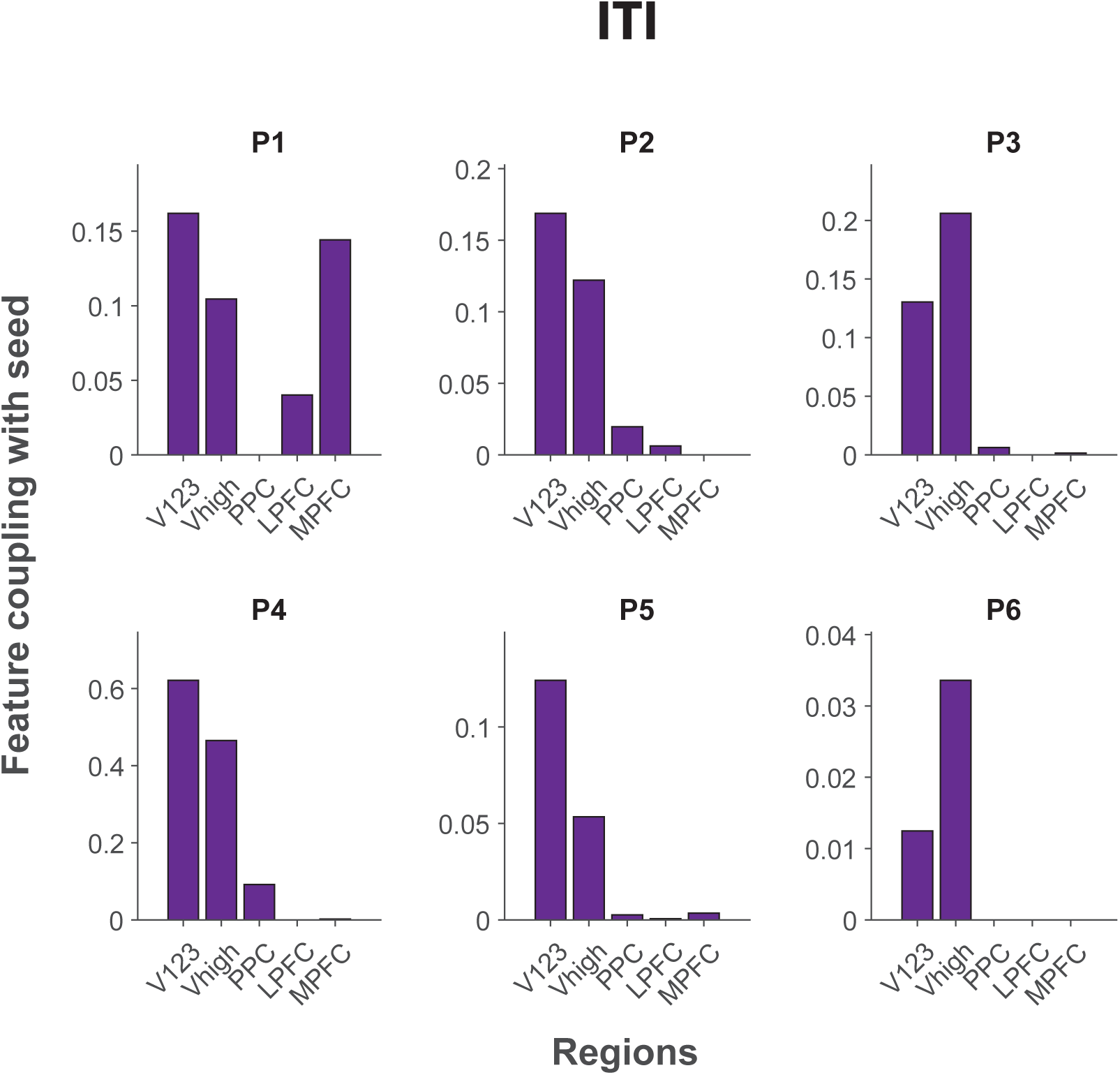
The same analysis as in Figure 7C, except here the epoch of the ITIs are taken as the psychological regressor, as opposed to the epoch of presentation of the visual stimuli In this situation, we did not observe robust coupling between mPFC value areas and lateral PFC and PPC, thereby supporting the possibility that increased coupling between lPFC, PPC and mPFC occurs specifically at the time of stimulus evaluation.

**Figure S13:**
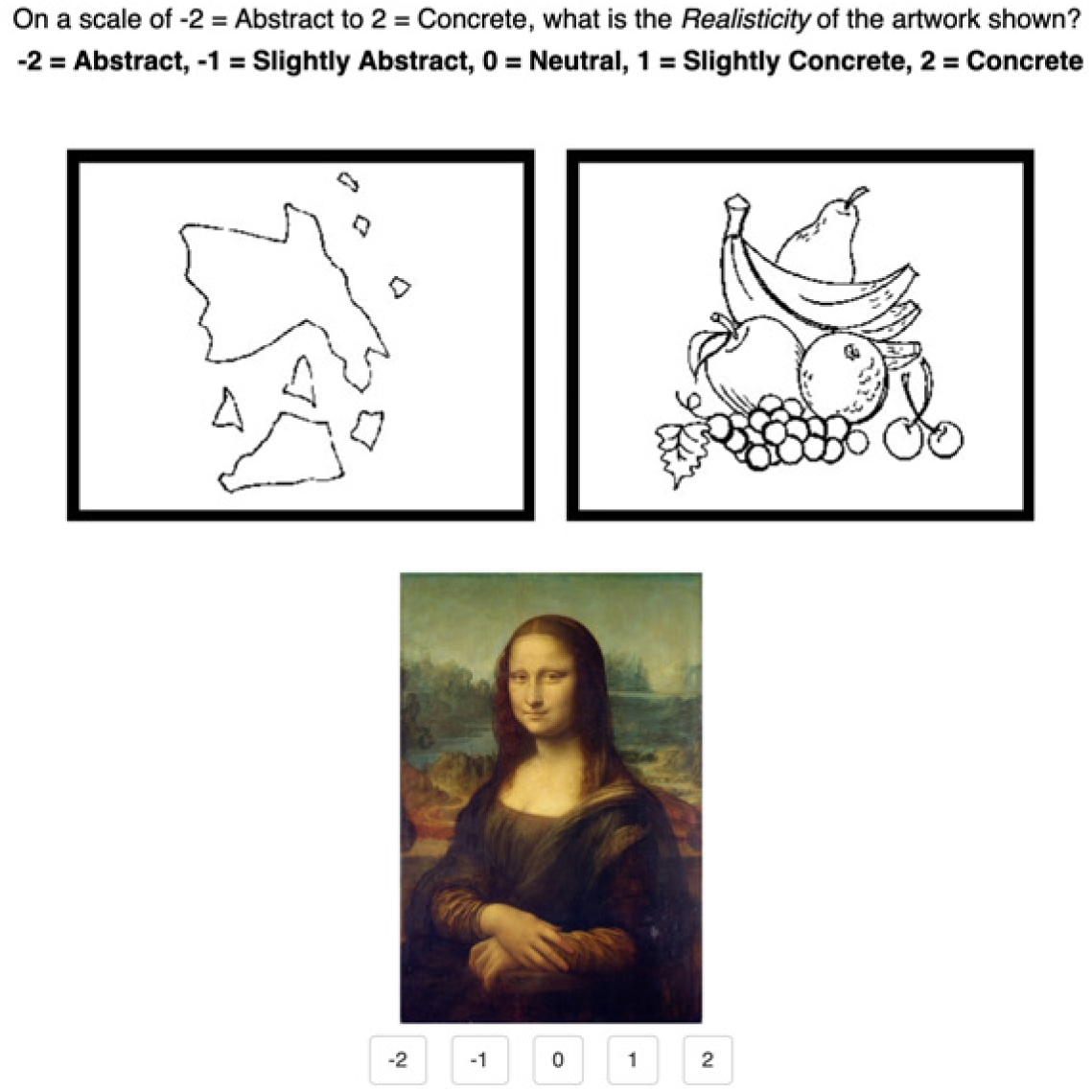
An example trial of feature annotation. Annotators were asked to evaluate high-level feature values (from −2 to 2), following.^26, 27^

**Figure S14:**
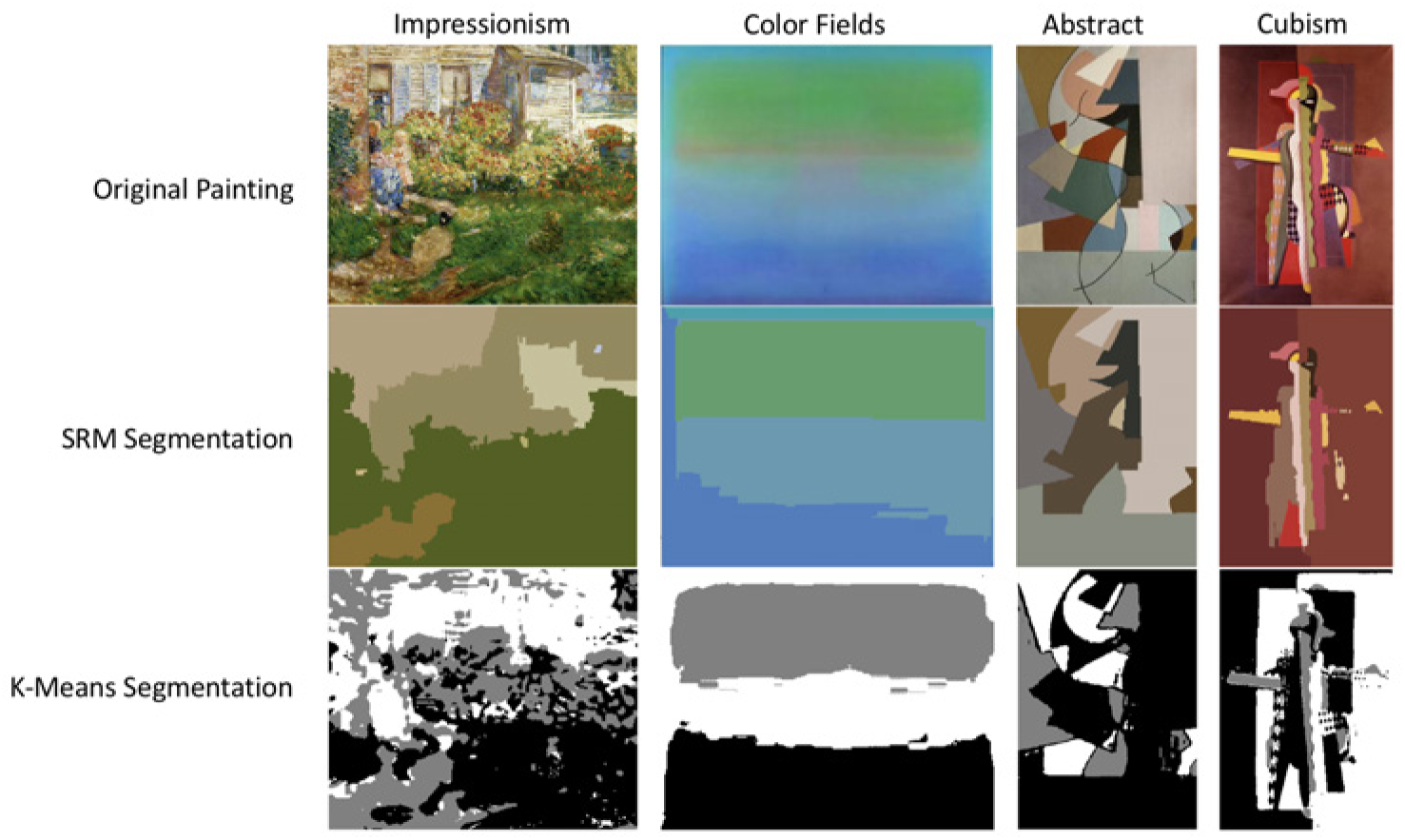
Statistical region merging (SRM) and k-means segmentation examples. Top: original images. Middle: SRM segmentation results. Bottom: k-means segmentation results.

## Footnotes

1 We thank Avi Vaidya and Lesley Fellows for this suggestion.

2 https://github.com/hiroyuki-kasai/SparseGDLibrary

## References

1. I. Kant, Critique of judgment. Hackett Publishing, 1987.

2. G. T. Fechner, Vorschule der aesthetik, vol. 1. Breitkopf & Hä rtel, 1876.

3. V. S. Ramachandran and W. Hirstein, “The science of art: A neurological theory of aesthetic experience,” Journal of consciousness Studies, vol. 6, no. 6-7, pp. 15–51, 1999.

4. S. Zeki, “Inner vision: An exploration of art and the brain,” 2002.

5. H. Leder, B. Belke, A. Oeberst, and D. Augustin, “A model of aesthetic appreciation and aesthetic judgments,” British journal of psychology, vol. 95, no. 4, pp. 489–508, 2004.

6. A. Chatterjee, “Neuroaesthetics: a coming of age story,” Journal of cognitive neuroscience, vol. 23, no. 1, pp. 53–62, 2011.

7. A. P. Shimamura and S. E. Palmer, Aesthetic science: Connecting minds, brains, and experience. OUP USA, 2012.

8. S. E. Palmer, K. B. Schloss, and J. Sammartino, “Visual aesthetics and human preference,” Annual review of psychology, vol. 64, pp. 77–107, 2013.

9. H. Leder and M. Nadal, “Ten years of a model of aesthetic appreciation and aesthetic judgments: The aesthetic episode–developments and challenges in empirical aesthetics,” British Journal of Psychology, vol. 105, no. 4, pp. 443–464, 2014.

10. A. Chatterjee, “Prospects for a cognitive neuroscience of visual aesthetics,” Bull. Psychol. Art, vol. 4, 2003.

11. C. J. Cela-Conde, G. Marty, F. Maestú, T. Ortiz, E. Munar, A. Ferná ndez, M. Roca, J. Rosselló, and F. Quesney, “Activation of the prefrontal cortex in the human visual aesthetic perception,” Proceedings of the National Academy of Sciences, vol. 101, no. 16, pp. 6321–6325, 2004.

12. H. Kawabata and S. Zeki, “Neural correlates of beauty,” Journal of neurophysiology, vol. 91, no. 4, pp. 1699–1705, 2004.

13. E. U. Weber and E. J. Johnson, “Constructing preferences from memory,” The Construction of Preference, Lichtenstein, S. & Slovic, P.,(eds.), pp. 397–410, 2006.

14. G. E. Wimmer and D. Shohamy, “Preference by association: how memory mechanisms in the hippocampus bias decisions,” Science, vol. 338, no. 6104, pp. 270–273, 2012.

15. H. C. Barron, R. J. Dolan, and T. E. Behrens, “Online evaluation of novel choices by simultaneous representation of multiple memories,” Nature neuroscience, vol. 16, no. 10, p. 1492, 2013.

16. C. M. Bishop, Pattern recognition and machine learning. springer, 2006.

17. S. Suzuki, L. Cross, and J. P. O’Doherty, “Elucidating the underlying components of food valuation in the human orbitofrontal cortex,” Nature neuroscience, vol. 20, no. 12, p. 1780, 2017.

18. J. D. Howard and J. A. Gottfried, “Configural and elemental coding of natural odor mixture components in the human brain,” Neuron, vol. 84, no. 4, pp. 857–869, 2014.

19. T. A. Hare, C. F. Camerer, and A. Rangel, “Self-control in decision-making involves modulation of the vmpfc valuation system,” Science, vol. 324, no. 5927, pp. 646–648, 2009.

20. S.-L. Lim, J. P. O’Doherty, and A. Rangel, “Stimulus value signals in ventromedial pfc reflect the integration of attribute value signals computed in fusiform gyrus and posterior superior temporal gyrus,” Journal of Neuroscience, vol. 33, no. 20, pp. 8729–8741, 2013.

21. T. Kahnt, J. Heinzle, S. Q. Park, and J.-D. Haynes, “Decoding different roles for vmpfc and dlpfc in multi-attribute decision making,” Neuroimage, vol. 56, no. 2, pp. 709–715, 2011.

22. V. Mante, D. Sussillo, K. V. Shenoy, and W. T. Newsome, “Context-dependent computation by recurrent dynamics in prefrontal cortex,” nature, vol. 503, no. 7474, p. 78, 2013.

23. G. Pelletier and L. K. Fellows, “A critical role for human ventromedial frontal lobe in value comparison of complex objects based on attribute configuration,” Journal of Neuroscience, vol. 39, no. 21, pp. 4124–4132, 2019.

24. S. J. Gershman, J. Malmaud, and J. B. Tenenbaum, “Structured representations of utility in combinatorial domains.,” Decision, vol. 4, no. 2, p. 67, 2017.

25. C. Li and T. Chen, “Aesthetic visual quality assessment of paintings,” IEEE Journal of selected topics in Signal Processing, vol. 3, no. 2, pp. 236–252, 2009.

26. A. Chatterjee, P. Widick, R. Sternschein, W. B. Smith, and B. Bromberger, “The assessment of art attributes,” Empirical Studies of the Arts, vol. 28, no. 2, pp. 207–222, 2010.

27. A. R. Vaidya, M. Sefranek, and L. K. Fellows, “Ventromedial frontal lobe damage alters how specific attributes are weighed in subjective valuation,” Cerebral Cortex, pp. 1–11, 2017.

28. C. Rother, V. Kolmogorov, and A. Blake, “Grabcut: Interactive foreground extraction using iterated graph cuts,” in ACM transactions on graphics (TOG), vol. 23, pp. 309–314, ACM, 2004.

29. N. Murray, L. Marchesotti, and F. Perronnin, “Ava: A large-scale database for aesthetic visual analysis,” in Computer Vision and Pattern Recognition (CVPR), 2012 IEEE Conference on, pp. 2408–2415, IEEE, 2012.

30. K. Simonyan and A. Zisserman, “Very deep convolutional networks for large-scale image recognition,” arXiv preprint arXiv:1409.1556, 2014.

31. J. Deng, W. Dong, R. Socher, L.-J. Li, K. Li, and L. Fei-Fei, “Imagenet: A large-scale hierarchical image database,” in 2009 IEEE conference on computer vision and pattern recognition, pp. 248–255, Ieee, 2009.

32. H. Hong, D. L. Yamins, N. J. Majaj, and J. J. DiCarlo, “Explicit information for category-orthogonal object properties increases along the ventral stream,” Nature neuroscience, vol. 19, no. 4, p. 613, 2016.

33. P. L. Smith and D. R. Little, “Small is beautiful: In defense of the small-n design,” Psychonomic bulletin & review, vol. 25, no. 6, pp. 2083–2101, 2018.

34. K. N. Kay, T. Naselaris, R. J. Prenger, and J. L. Gallant, “Identifying natural images from human brain activity,” Nature, vol. 452, no. 7185, p. 352, 2008.

35. C. Padoa-Schioppa and J. A. Assad, “Neurons in the orbitofrontal cortex encode economic value,” Nature, vol. 441, no. 7090, p. 223, 2006.

36. J. W. Kable and P. W. Glimcher, “The neural correlates of subjective value during intertemporal choice,” Nature neuroscience, vol. 10, no. 12, p. 1625, 2007.

37. J. Gläscher, A. N. Hampton, and J. P. O’Doherty, “Determining a role for ventromedial prefrontal cortex in encoding action-based value signals during reward-related decision making,” Cerebral cortex, vol. 19, no. 2, pp. 483–495, 2008.

38. F. Grabenhorst and E. T. Rolls, “Value, pleasure and choice in the ventral prefrontal cortex,” Trends in cognitive sciences, vol. 15, no. 2, pp. 56–67, 2011.

39. T. Ishizu and S. Zeki, “The brain’s specialized systems for aesthetic and perceptual judgment,” European Journal of Neuroscience, vol. 37, no. 9, pp. 1413–1420, 2013.

40. L. Wang, R. E. Mruczek, M. J. Arcaro, and S. Kastner, “Probabilistic maps of visual topography in human cortex,” Cerebral cortex, vol. 25, no. 10, pp. 3911–3931, 2014.

41. L. Wang, R. E. Mruczek, M. J. Arcaro, and S. Kastner, “Probabilistic maps of visual topography in human cortex,” Cerebral cortex, vol. 25, no. 10, pp. 3911–3931, 2014.

42. J. S. Baizer, L. G. Ungerleider, and R. Desimone, “Organization of visual inputs to the inferior temporal and posterior parietal cortex in macaques,” Journal of Neuroscience, vol. 11, no. 1, pp. 168–190, 1991.

43. S. C. Rao, G. Rainer, and E. K. Miller, “Integration of what and where in the primate prefrontal cortex,” Science, vol. 276, no. 5313, pp. 821–824, 1997.

44. M. Rigotti, O. Barak, M. R. Warden, X.-J. Wang, N. D. Daw, E. K. Miller, and S. Fusi, “The importance of mixed selectivity in complex cognitive tasks,” Nature, vol. 497, no. 7451, p. 585, 2013.

45. C. Y. Zhang, T. Aflalo, B. Revechkis, E. R. Rosario, D. Ouellette, N. Pouratian, and R. A. Andersen, “Partially mixed selectivity in human posterior parietal association cortex,” Neuron, vol. 95, no. 3, pp. 697–708, 2017.

46. M. Noonan, M. Walton, T. Behrens, J. Sallet, M. Buckley, and M. Rushworth, “Separate value comparison and learning mechanisms in macaque medial and lateral orbitofrontal cortex,” Proceedings of the National Academy of Sciences, vol. 107, no. 47, pp. 20547– 20552, 2010.

47. E. A. Vessel, G. G. Starr, and N. Rubin, “The brain on art: intense aesthetic experience activates the default mode network,” Frontiers in human neuroscience, vol. 6, p. 66, 2012.

48. Y. Bengio, “Deep learning of representations for unsupervised and transfer learning,” in Proceedings of ICML workshop on unsupervised and transfer learning, pp. 17–36, 2012.

49. M. Leshno, V. Y. Lin, A. Pinkus, and S. Schocken, “Multilayer feedforward networks with a nonpolynomial activation function can approximate any function,” Neural networks, vol. 6, no. 6, pp. 861–867, 1993.

50. T. Hofmann, B. Schö lkopf, and A. J. Smola, “Kernel methods in machine learning,” The annals of statistics, pp. 1171–1220, 2008.

51. D. C. Van Essen and J. H. Maunsell, “Hierarchical organization and functional streams in the visual cortex,” Trends in neurosciences, vol. 6, pp. 370–375, 1983.

52. D. J. Felleman and D. E. Van, “Distributed hierarchical processing in the primate cerebral cortex.,” Cerebral cortex (New York, NY: 1991), vol. 1, no. 1, pp. 1–47, 1991.

53. S. Hochstein and M. Ahissar, “View from the top: Hierarchies and reverse hierarchies in the visual system,” Neuron, vol. 36, no. 5, pp. 791–804, 2002.

54. C. S. Konen and S. Kastner, “Two hierarchically organized neural systems for object information in human visual cortex,” Nature neuroscience, vol. 11, no. 2, p. 224, 2008.

55. C. F. Cadieu, H. Hong, D. L. Yamins, N. Pinto, D. Ardila, E. A. Solomon, N. J. Majaj, and J. J. DiCarlo, “Deep neural networks rival the representation of primate it cortex for core visual object recognition,” PLoS computational biology, vol. 10, no. 12, p. e1003963, 2014.

56. S.-M. Khaligh-Razavi and N. Kriegeskorte, “Deep supervised, but not unsupervised, models may explain it cortical representation,” PLoS computational biology, vol. 10, no. 11, p. e1003915, 2014.

57. U. Güçlü and M. A. van Gerven, “Deep neural networks reveal a gradient in the complexity of neural representations across the ventral stream,” Journal of Neuroscience, vol. 35, no. 27, pp. 10005–10014, 2015.

58. D. Brieber, M. Nadal, and H. Leder, “In the white cube: Museum context enhances the valuation and memory of art,” Acta psychologica, vol. 154, pp. 36–42, 2015.

59. O. Esteban, R. Blair, C. J. Markiewicz, S. L. Berleant, C. Moodie, F. Ma, A. I. Isik, A. Erramuzpe, M. Kent, James D. and Goncalves, E. DuPre, K. R. Sitek, D. E. P. Gomez, D. J. Lurie, Z. Ye, R. A. Poldrack, and K. J. Gorgolewski, “fmriprep,” Software, 2018.

60. K. J. Gorgolewski, O. Esteban, C. J. Markiewicz, E. Ziegler, D. G. Ellis, M. P. Notter, D. Jarecka, H. Johnson, C. Burns, A. Manhã es-Savio, C. Hamalainen, B. Yvernault, T. Salo, K. Jordan, M. Goncalves, M. Waskom, D. Clark, J. Wong, F. Loney, M. Modat, B. E. Dewey, C. Madison, M. Visconti di Oleggio Castello, M. G. Clark, M. Dayan, D. Clark, A. Keshavan, B. Pinsard, A. Gramfort, S. Berleant, D. M. Nielson, S. Bougacha, G. Varoquaux, B. Cipollini, R. Markello, A. Rokem, B. Moloney, Y. O. Halchenko, D. Wassermann, M. Hanke, C. Horea, J. Kaczmarzyk, G. de Hollander, E. DuPre, A. Gillman, D. Mordom, C. Buchanan, R. Tungaraza, W. M. Pauli, S. Iqbal, S. Sikka, M. Mancini, Y. Schwartz, I. B. Malone, M. Dubois, C. Frohlich, D. Welch, J. Forbes, J. Kent, A. Watanabe, C. Cumba, J. M. Huntenburg, E. Kastman, B. N. Nichols, A. Eshaghi, D. Ginsburg, A. Schaefer, B. Acland, S. Giavasis, J. Kleesiek, D. Erickson, R. Kü ttner, C. Haselgrove, C. Correa, A. Ghayoor, F. Liem, J. Millman, D. Haehn, J. Lai, D. Zhou, R. Blair, T. Glatard, M. Renfro, S. Liu, A. E. Kahn, F. Pé rez-Garćıa, W. Triplett, L. Lampe, J. Stadler, X.-Z. Kong, M. Hallquist, A. Chetverikov, J. Salvatore, A. Park, R. Poldrack, R. C. Craddock, S. Inati, O. Hinds, G. Cooper, L. N. Perkins, A. Marina, A. Mattfeld, M. Noel, L. Snoek, K. Matsubara, B. Cheung, S. Rothmei, S. Urchs, J. Durnez, F. Mertz, D. Geisler, A. Floren, S. Gerhard, P. Sharp, M. Molina-Romero, A. Weinstein, W. Broderick, V. Saase, S. K. Andberg, R. Harms, K. Schlamp, J. Arias, D. Papadopoulos Orfanos, C. Tarbert, A. Tambini, A. De La Vega, T. Nickson, M. Brett, M. Falkiewicz, K. Podranski, J. Linkersdö rfer, G. Flandin, E. Ort, D. Shachnev, D. McNamee, A. Davison, J. Varada, I. Schwabacher, J. Pellman, M. Perez-Guevara, R. Khanuja, N. Pannetier, C. McDermottroe, and S. Ghosh, “Nipype,” Software, 2018.

61. N. J. Tustison, B. B. Avants, P. A. Cook, Y. Zheng, A. Egan, P. A. Yushkevich, and J. C. Gee, “N4itk: Improved n3 bias correction,” IEEE Transactions on Medical Imaging, vol. 29, no. 6, pp. 1310–1320, 2010.

62. B. Avants, C. Epstein, M. Grossman, and J. Gee, “Symmetric diffeomorphic image registration with cross-correlation: Evaluating automated labeling of elderly and neurodegenerative brain,” Medical Image Analysis, vol. 12, no. 1, pp. 26–41, 2008.

63. Y. Ke, X. Tang, and F. Jing, “The design of high-level features for photo quality assessment,” in 2006 IEEE Computer Society Conference on Computer Vision and Pattern Recognition (CVPR’06), vol. 1, pp. 419–426, IEEE, 2006.

64. M. B. Salah, A. Mitiche, and I. B. Ayed, “Multiregion image segmentation by parametric kernel graph cuts,” IEEE Transactions on Image Processing, vol. 20, no. 2, pp. 545–557, 2010.

65. R. Nock and F. Nielsen, “Statistical region merging,” IEEE Transactions on pattern analysis and machine intelligence, vol. 26, no. 11, pp. 1452–1458, 2004.

66. Q. Zhu, M.-C. Yeh, K.-T. Cheng, and S. Avidan, “Fast human detection using a cascade of histograms of oriented gradients,” in 2006 IEEE Computer Society Conference on Computer Vision and Pattern Recognition (CVPR’06), vol. 2, pp. 1491–1498, IEEE, 2006.

67. M. Hein and T. Bühler, “An inverse power method for nonlinear eigenproblems with applications in 1-spectral clustering and sparse pca,” in Advances in Neural Information Processing Systems, pp. 847–855, 2010.

68. N. Murray and A. Gordo, “A deep architecture for unified aesthetic prediction,” arXiv preprint arXiv:1708.04890, 2017.

69. P. J. Huber, “Robust estimation of a location parameter,” Ann. Math. Statist., vol. 35, pp. 73–101, 03 1964.

70. L. A. Gatys, A. S. Ecker, and M. Bethge, “Image style transfer using convolutional neural networks,” in Proceedings of the IEEE conference on computer vision and pattern recognition, pp. 2414–2423, 2016.

